# World Wide Eyes: Social Media Assisted Wildlife Research. Assessing Supplementary Feeding Areas for Eastern Australian Humpback Whales as a Case Study

**DOI:** 10.64898/2026.10.26.761931

**Authors:** G. Marzorati, E. Valsecchi

## Abstract

Humpback whales typically adopt a “feast-and-famine” strategy, feeding intensively at high-latitude summer feeding grounds while fasting throughout migration. However, increasing reports of feeding at mid-latitudes suggest that some individuals may supplement their energetic requirements along migratory corridors. Detecting and monitoring these emerging behaviours remains challenging because conventional marine mammal surveys are often spatially and financially constrained. Here, we used passive citizen science by systematically compiling humpback whale feeding records from publicly shared Facebook and Instagram posts and the citizen-science platform Happywhale to investigate feeding events of the Eastern Australian population over five consecutive migration seasons (2020– 2024). A total of 256 feeding records were collected, revealing three recurrent candidate Supplementary Feeding Areas (SFAs): the Sapphire Coast (New South Wales), Port Phillip and Bass Strait (Victoria), and southeast Tasmania. Feeding events occurred across multiple migratory months, involved both solitary whales and groups, and included both krill and schooling fish as prey, with lunge feeding representing the predominant feeding strategy. Comparison of passive and active citizen-science sources showed that social-media records substantially increased the number of documented feeding events and more frequently allowed feeding behaviour to be identified. The temporal recurrence of feeding events across five consecutive migration seasons supports the hypothesis that some behaviours traditionally interpreted as opportunistic feeding may instead represent recurrent supplementary feeding. Overall, this study demonstrates the value of passive citizen science as a cost-effective complement to conventional monitoring programmes for detecting large-scale behavioural changes and identifying candidate supplementary feeding areas in migratory humpback whales.

## INTRODUCTION

Humpback whale populations undertake seasonal migration from winter’s low-latitude breeding grounds to summer’s high-latitude feeding grounds (Stockin et al. 2005; Andrews-Goff et al. 2018). The Eastern Australian humpback whale population is designated as E1 breeding stock, as identified by the IWC (Gales et al. 2009). The E1 breeding stock feeds in the circumpolar waters around Antarctica (Area V: 120° - 180° E) before then migrating along the Eastern coast of Australia to breed in the waters circumscribed by the Great Barrier Reef (16-21° S) (Chittleborough 1965; Simmons and Marsh 1986; Franklin et al. 2017; Bettridge et al. 2015; see also Figure 1 in Andrews-Goff et al. 2018).

**Figure 1.**
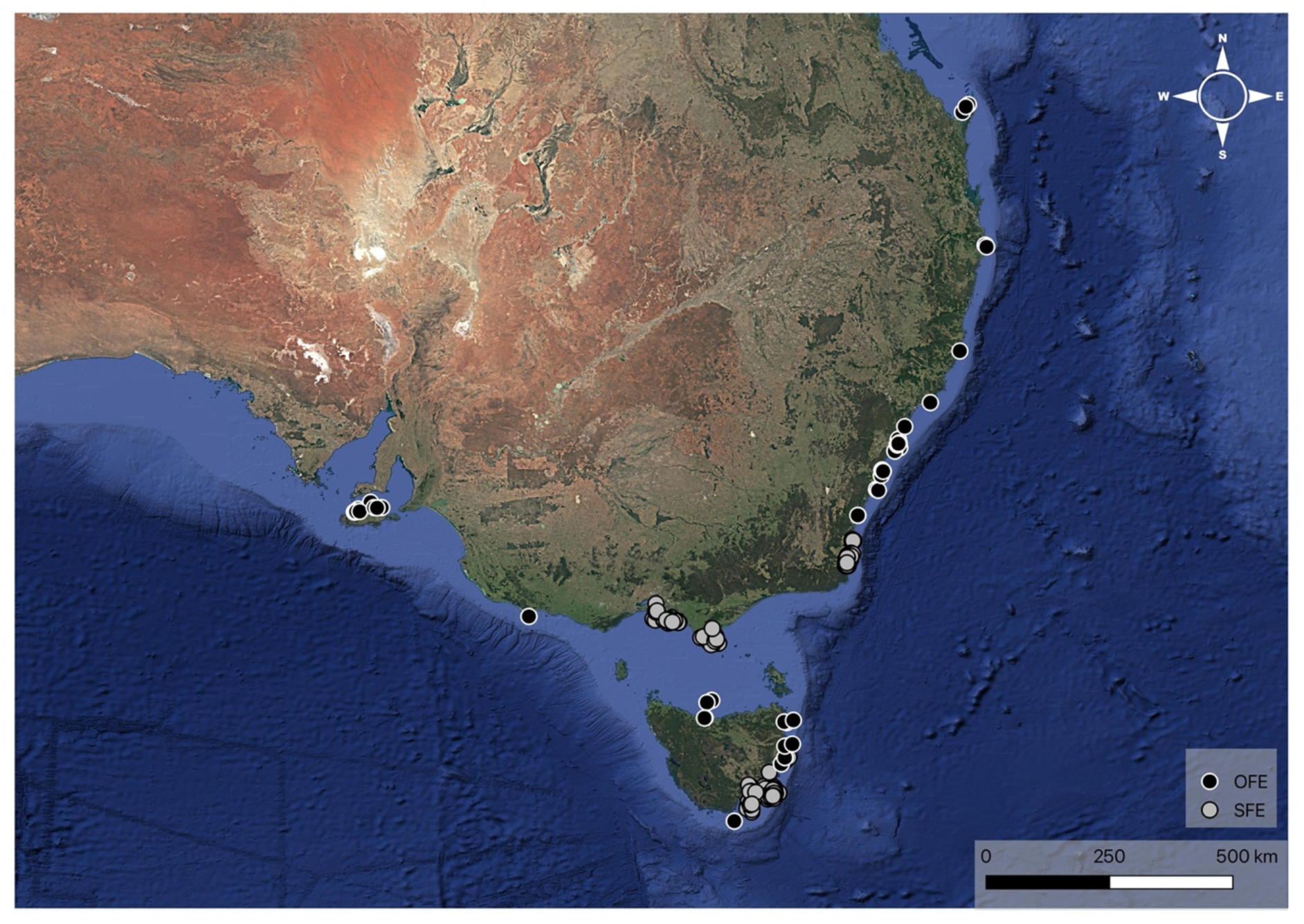
Map of the humpback whales’ feeding events sightings in 2020, 2021, 2022, 2023 and 2024 migratory seasons. Opportunistic feeding events (OFE) are shown in black, while those events fulfilling the criteria for being considered as supplementary feeding events (SFE) are shown in light grey.

Humpback whales are known to adopt a particular feeding regime shaped around the consumer– resource interactions characterizing their life-cycle: the “feast and famine” strategy (Andrews-Goff et al. 2018). They feed at high-latitude feeding grounds, stocking up all the energy needed for the entire year while they fast during the migration (while travelling to and from the breeding grounds) and while they remain on the breeding ground (Dawbin 1966; Craig et al., 2003; Groß et al. 2020). The consequence of this approach is that they rely only on the energy reserves accumulated in the feeding grounds during the summer for migrating, breeding, calving, and nursing. Hence, successful migration and reproduction are assumed to directly depend on the stored energy and the availability of food resources on the feeding grounds (Groß et al. 2020; Alerstam et al. 2003; Craig et al. 2003; Owen et al. 2015). Foraging methods and prey type vary according to the different physical and chemical characteristics of the various geographic areas and the size and amount of prey present (Friedlaender et al. 2009; Hays et al. 1985; Garcia et al. 2021).

Considering the amount of energy consumed during the migration (Craig et al. 2003), migratory whales should optimize their intake of high-quality prey during their summer feedings (Witteveen et al. 2011), but additionally, they could also opportunistically take advantage of ephemeral food sources when available in considerable quantity (Silva et al. 2013). More recently, researchers hypothesized that supplementary feeding may occur when energetic requirements are unmet on Antarctic feeding grounds (Andrews-Goff et al. 2018).

In support of this hypothesis, there are few sightings reported in the past few years of feeding humpback whales away from the feeding grounds such as in Brazil (Danilewicz et al. 2009), St. Mary Bay (Nova Scotia, Canada) (Askin et al. 2017) and across the southern Benguela System (along the coast of South Africa between St. Helena Bay and Cape Point) (Findlay et al. 2017). Global climate change is expected to significantly affect the productivity of circumpolar, and thus also Antarctic, waters, with rising temperatures leading to shifts in ocean circulation patterns and altered nutrient availability, which could have cascading effects on marine ecosystems (Vaughan et al. 2014; Stroeve et al. 2007; Stroeve et al. 2018). In future years, continued warming is likely to reduce primary productivity in some regions, while in others, increased nutrient input could stimulate short-term productivity changes, ultimately threatening the stability of the ecosystem (Clarke et al., 2007). Humpback whales may gradually adapt their feeding habits to climate change, with supplementary feeding becoming increasingly important. Feeding in East Australia has been observed more regularly than elsewhere (Kaufman et al. 2006; Silva et al. 2013). Due to recent observations and satellite tracks of the migration, scientists noticed that E1 humpback whales have multiple feeding stops lasting days to weeks in several highly productive temperate areas along the migration route (Groß et al. 2020). Even analyses on stable isotopes of baleen whale plates indicate that supplemental feeding may be quite a common strategy for East Australian humpback whales (Andrews-Goff et al. 2018; Eisenmann et al. 2016). Additional data come from the studies of Gill et al. 1998, Gedamke et al. 2007, Stamation et al. 2007, Gales et al. 2009; Silva et al. 2013, Findlay et al. 2017, Owen et al. 2017 and Pirotta et al. 2021. However, information on supplementary feeding for E1 humpback whales is still patchy, being affected by an uneven distribution of research efforts.

To address these limitations, traditional marine mammal monitoring programs remain logistically demanding, geographically restricted and financially expensive, limiting their capacity to detect rapidly emerging ecological responses to environmental change. As climate-driven shifts in ocean productivity increasingly alter prey distributions and availability, there is a growing need for monitoring approaches capable of providing continuous, broad-scale observations across large spatial extents.

Within this context, citizen science (CS), or participatory science, is experiencing exponential growth in countless scientific fields and, in particular, in conservation biology (Follett et al. 2015; MacPhail and Colla 2020). For instance, in the international network Global Biodiversity Information Facility (www.gbif.org), 50% of occurrence records are Citizen Science observations and 6 of the top 10 datasets are citizen science datasets (Waller 2019). This increase is due to multiple reasons, ranging from greater awareness and desire to take part in conservation, on the part of the public (Lewandowski et al. 2017; MacPhail et al. 2020; Maund et al. 2020), and the need to have a task force spread across large territories, on the researchers’ part (Van Der Velde et al. 2017; Horns et al. 2018; MacPhail and Colla 2020). However, in most CS projects, participants “actively” choose to take part in the research, and courses or induction briefings are often provided prior to data collection (Edwards et al. 2021).

There are also alternative forms of CS that are less explored, such as those defined as “passive” (Edwards et al. 2021), since data are collected unintentionally and made spontaneously public by social-media users, unaware of their scientific value. An emblematic example is when scientifically useful data are extrapolated from videos and photographs that digital community members spontaneously publish on social media (Sullivan et al. 2019; Ghermandi et al. 2019; Edwards et al. 2021; Morais et al. 2021). According to the metrics associated with the publication of videos on social media, it appears that the drive making the average social platforms’ user willing to post a video is the impression of having filmed a video that is (characteristics listed in order of importance according to the metrics provided by the site www.vidjet.com): funny, creative, informative, relaxing, inspirational, engaging, trendy, exciting, relevant, helpful/how-to. Thus, about wildlife filming, the user might be prone to publish anomalous situations, animals considered dangerous (such as sharks and snakes), or animals difficult to observe and rare, or charismatic. Whales certainly fall into these last two categories, and the flourishing of the whale-watching industry demonstrates the appeal these animals (and their relatives, dolphins, and seals) arouse in the general public. Passive citizen science therefore represents a potentially cost-effective and accessible complement to conventional survey methods (Lodi & Tardin 2018; Cranswick et al. 2022). By exploiting existing observational material rather than relying on dedicated sampling effort, it may facilitate the detection of rare behaviours, emerging ecological phenomena, and conservation-relevant changes that would otherwise remain unrecorded (Morais et al. 2021; Cranswick et al. 2022).

This study aims to assess the effectiveness of a “passive” citizen science approach, based on the analysis of videos and photos published on community media to provide additional evidence about supplementary feeding events in humpback whales migrating off Eastern Australia, relying on data collected in areas where this behavior has been barely investigated.

## MATERIALS AND METHODS

The study was carried out by retrieving CS data collected from April 2020 to January 2025, focusing on the migratory corridor of the E1 Humpback whale population, spanning from the east coast of Australia to Tasmania. Because humpback whale migration seasons extend from April/May to January of the following year, we analysed five consecutive migration seasons across Eastern Australia. Consequently, records collected in January were assigned to the migration season of the previous calendar year; for example, data collected in January 2025 were included in the 2024 migration season. Following this seasonal framework, records collected from April to August were classified as the northern migration, corresponding to the movement of E1 humpback whales from their Antarctic feeding grounds to the breeding areas around Hervey Bay, whereas records collected from September to January were classified as the southern migration, representing the return migration from the breeding grounds to the Antarctic feeding areas.

### Data Collection

The data of this study were gathered through passive citizen science. The sightings were collected from posts on social media and the public platforms Facebook and Instagram, and from a website called Happywhale (www.happywhale.com), a platform engaging citizen scientists to identify individual marine mammals by collecting and submitting photo-identification records (Cheeseman et al. 2017), thus documenting whale sightings worldwide. Facebook and Instagram were the two social media platforms selected for this study, based on two primary criteria: they rank among the most widely used platforms globally, making them well-suited for collecting a sufficiently large dataset to ensure statistical robustness; and they appeal to a relatively broad demographic spectrum, cutting across different age groups. Moreover, Facebook and Instagram are the two social media platforms most consistently identified in the conservation literature as suitable sources of georeferenced user-generated content for biodiversity and wildlife monitoring, owing to the availability of photographs, videos, hashtags, place tags, and searchable metadata (Di Minin et al., 2015; Toivonen et al., 2019; Lenzi et al., 2020; Edwards et al., 2021).

The research employed a detailed methodology using keyword-based searches across social media platforms and the HappyWhale website to document humpback whale feeding behaviors in Australian waters. Keywords used on Instagram and Facebook to locate posts, photos, and videos were: “humpback whale,” “humpback whale feeding,” “humpback whale drone,” “humpback whale Australia,” “humpback whale eat,” and “humpback whale eating”. Profiles of frequent contributors, such as whale-watching companies and drone photographers, were further analyzed for overlooked content, given their expertise and access to high-quality equipment. Data with an unclear date or not specified location were not considered.

Occasionally, the search also retrieved Facebook or Instagram posts sharing online news articles reporting humpback whale feeding events. These records were included in the passive citizen science dataset because they were retrieved through social media. However, for the comparison between passive and active citizen science sources, they were treated as a separate category to distinguish journalistic content from user-generated social media posts.

On Facebook, marine mammal sighting groups were examined, and posts were cross-verified through comments to confirm feeding behaviors, though location and time information were sometimes limited. The HappyWhale platform was also utilized to search for humpback whale sightings in Australian migration zones (E1 and D stocks), filtering results by species, location, and time frame. Descriptions were analyzed for feeding activity details, and any gaps in the data were cross-referenced with social media findings. This integrated approach combined user-generated content and professional observations to compile the final dataset

Only records of feeding E1 humpback whales collected along the east coast of Australia between April 2020 and January 2025 were included in the analysis. To be considered, each record had to contain complete information on the sighting date, location, and pod size, either in the original post description or in the associated comments. For records from whale-watching companies, the sighting location was inferred from the operator’s known area of activity or, when available, from the location information provided in the company profile. Records lacking any of these three key variables or containing highly uncertain information were excluded from the analysis.

Potential duplicate records were identified as sightings reporting the same pod at the same location on the same date, with a comparable number of individuals. Records sharing the same date and reported locality were subsequently examined individually using all available contextual information, such as comments or source, to determine whether they could be confidently attributed to the same feeding event. Only records meeting these criteria would have been merged into a single observation for the analyses, while any complementary information provided by additional reports would have been retained.

Exact geographic coordinates were directly available only for records obtained from Happywhale. For records collected from Instagram and Facebook, coordinates were inferred from the location indicated in the post, its description, or the associated comments. Similarly, the time of the sighting, when available, was extracted from the post description or comments across all platforms.

When available, any additional information about the feeding strategy or the type of prey was recorded and analyzed to identify eventual preferences related to feeding activities. In a few cases, photos and videos were detailed enough to allow adding further information on the feeding technique and prey items.

Although Happywhale allows the photo-identification of individual humpback whales, individual re-identification was not included among the objectives of the present study. Consequently, no systematic matching of individuals across years or locations was performed, and all analyses were conducted at the feeding-event level rather than the individual level.

Passive citizen science provides non-standardised observational data that may be subject to spatial and temporal reporting biases. These limitations were considered during data interpretation. To further assess potential spatial differences in observation opportunity, whale-watching operator distribution was used as a qualitative proxy of observation effort (Supplementary Table S2). The number of whale-watching operators operating within each predefined region was compiled from publicly available information and compared with the distribution of feeding records. Regions were classified according to the number of operators as Low (1–2 operators), Moderate (3–4 operators), High (5–6 operators), and Very High (≥7 operators). This proxy was not used to correct or normalize feeding records, but only to provide contextual information on potential spatial variation in observation effort (Supplementary Table S3).

### Definition and annotation of Supplementary Feeding Areas

Supplementary feeding is considered a feeding action that occurs beyond the regular summer high-latitude feeding grounds and aims to supplement the energy lost during migration (Seyboth et al. 2023). It is arduous to distinguish supplementary feeding events from opportunistic feeding ones because they both occur during the migration. Still, opportunistic feeding is the random act when a whale takes advantage of a high prey concentration; conversely, supplementary feeding is the voluntary act of feeding to supplement the energy lost during the migration. These two behaviours should not necessarily be viewed as mutually exclusive. Rather, supplementary feeding may develop through the repeated exploitation of predictable prey aggregations initially encountered opportunistically. If prey resources remain consistently available across years, whales may repeatedly return to the same areas during migration, progressively incorporating these locations into their migratory foraging strategy. Under this framework, supplementary feeding can be interpreted as the long-term expression of recurrent opportunistic feeding events in predictable and profitable habitats. Based on this conceptual framework, this study considered candidate supplementary feeding behavior, a repeated feeding activity documented within the same geographic area over five consecutive migration seasons.

Due to the recent recognition of the phenomenon of supplementary feeding for humpback whales, neither a unanimous definition nor a quantitave criterion currently exist to identify a supplementary feeding area (SFA). In this study, a putative supplementary feeding area was considered a region with more than 50 feeding events from 2020 to 2024, with feeding activity documented in each of the five years. The threshold of 50 feeding events was established as an operational rather than a biological criterion. It was based on a minimum of ten feeding events per year, equivalent to approximately one documented feeding event per day over ten days each migration season, sustained across all five study years without any year lacking feeding records. This relatively high threshold was intentionally adopted to distinguish recurrent feeding behaviour from isolated opportunistic feeding events. Although the numerical value is necessarily arbitrary, it was chosen to represent a level of temporal persistence unlikely to result from chance observations alone.

This approach is consistent with the conceptual framework adopted in this study, whereby supplementary feeding is interpreted as the repeated use of predictable feeding habitats over multiple migration seasons rather than as isolated opportunistic feeding events. Because the behavioural intention of individual whales cannot be directly assessed, recurrence in both space and time provides the most objective criterion currently available for distinguishing these two phenomena.

The ten events considered in this study as the threshold for evaluating possible supplementary feeding grounds were usually not on consecutive days or in close temporal proximity. An exception is made for the “megapods” phenomenon, where large whale-pods were recorded on eight consecutive days (6-13/09/2021) at Bermagui (Sapphire Coast) where these events were considered as one per day. To note that “megapod” is a word used descriptively to indicate exceptionally large feeding aggregations comprising several tens to hundreds of humpback whales feeding simultaneously.

The map was created using QGIS. None of the observers took precise coordinates of the feeding events; consequently, the locations are an estimation based on the position delineated in the posts. Locating the various feeding events on a chart, some actions were visibly isolated from the others and were, therefore, considered opportunistic feeding events. In contrast, others were gathered in specific sites and repeated throughout the years (considered possible supplementary feeding events).

### Prey item analysis

When the recovered material allowed the prey item to be identified, further analyses were carried out to define whether the standard foraging method and prey target, studied on the E1 population in feeding areas in Antarctica, were maintained even at lower latitudes in putative supplementary feeding areas.

### Feeding behaviour

When it was possible to assess feeding-whales’ group size or to infer if whales were feeding alone or within pods, this behavior was noted, and, to assess whether the relationship between singletons and pods differs significantly between the Northern and Southern migrations, a chi-square (χ²) test was conducted with a significance level of α = 0.05.

## RESULTS

A total of 256 feeding records were collected. A detailed list of all feeding sightings is shown in Supplementary Table 1, where date, time, location, number of whales, and source are reported. Out of the 256 records, 60 (23.4%) were retrieved from Happywhale, while the remaining 196 (76.6%) were recovered from Instagram and Facebook. Considering that E1 humpback whale stock annually migrates across Australia from late April-beginning of May till December, and in order to investigate whether migrating animals make use of supplementary feeding, our analysis focused at first on this time window, which, in fact, accounted for the majority of sightings (n=251 out of 256, 98.05 %). However, the remaining five (5 out of 256, 1.95%) sightings occurring outside this eight-month window all fell within the first few weeks of January (no record was reported after the 16^th^ of January -precisely in year 2025). No record was reported in February, March, and April (with one exception, in Tasmania in 2020, when the first observation of a feeding northbound whale was recorded on the 23^rd^ of April). Thus, those five feeding-activities observed in January over the five years refer to whales that should be considered as late southbound migrants, rather than non-migrating whales.

The map in Figure 1 shows the locations of the different sightings reported in Eastern Australia and Tasmania. Three specific zones stood out for having more than 50 events of feeding humpback whales spread over the five years analysed (2020, 2021, 2022, 2023, 2024). In these three areas occurred the majority (n=193 out of 256, 75.4%) of 256 feeding events recorded in the five years from May to mid-January. To evaluate whether this spatial pattern could be influenced by differences in observation opportunity, the number of whale-watching operators was compared among regions using a qualitative observation effort proxy (Supplementary Table S3). These areas are: 1) the Sapphire Coast on the southern Coast of New South Wales, from Narooma to Eden (Twofold Bay), roughly extending between −36.2° and −37.1° in latitude; 2) Port Phillip and Bass Strait in Victoria, including waters surrounding Phillip Island up to the Wilsons Promontory (Victoria), spanning between −38.3° and −39.2° in latitude; 3) the Eastern coast of Tasmania, from Hobart, inside the river Derwent up to its mouth opening on the Tasman Sea, till South Bruny Island extending in width to the Eastern coast of the Tasman Peninsula (at the level of Pirates Bay and Eaglehawk Neck), spanning in latitude between −42.9° and −43.4°.

The total number of sightings of feeding humpback whales collected throughout the migratory season of 2020, 2021, 2022, 2023, and 2024 was 87 in Tasmania, 50 in Port Phillip/Bass Strait, and 56 in the Sapphire Coast. Notably, the regions with the highest number of feeding records did not correspond to those with the highest whale-watching operator presence, suggesting that spatial differences in records were not solely explained by observation opportunity. Data for the overall dataset and for the five study years and areas are summarized in Figure 2.

**Figure 2.**
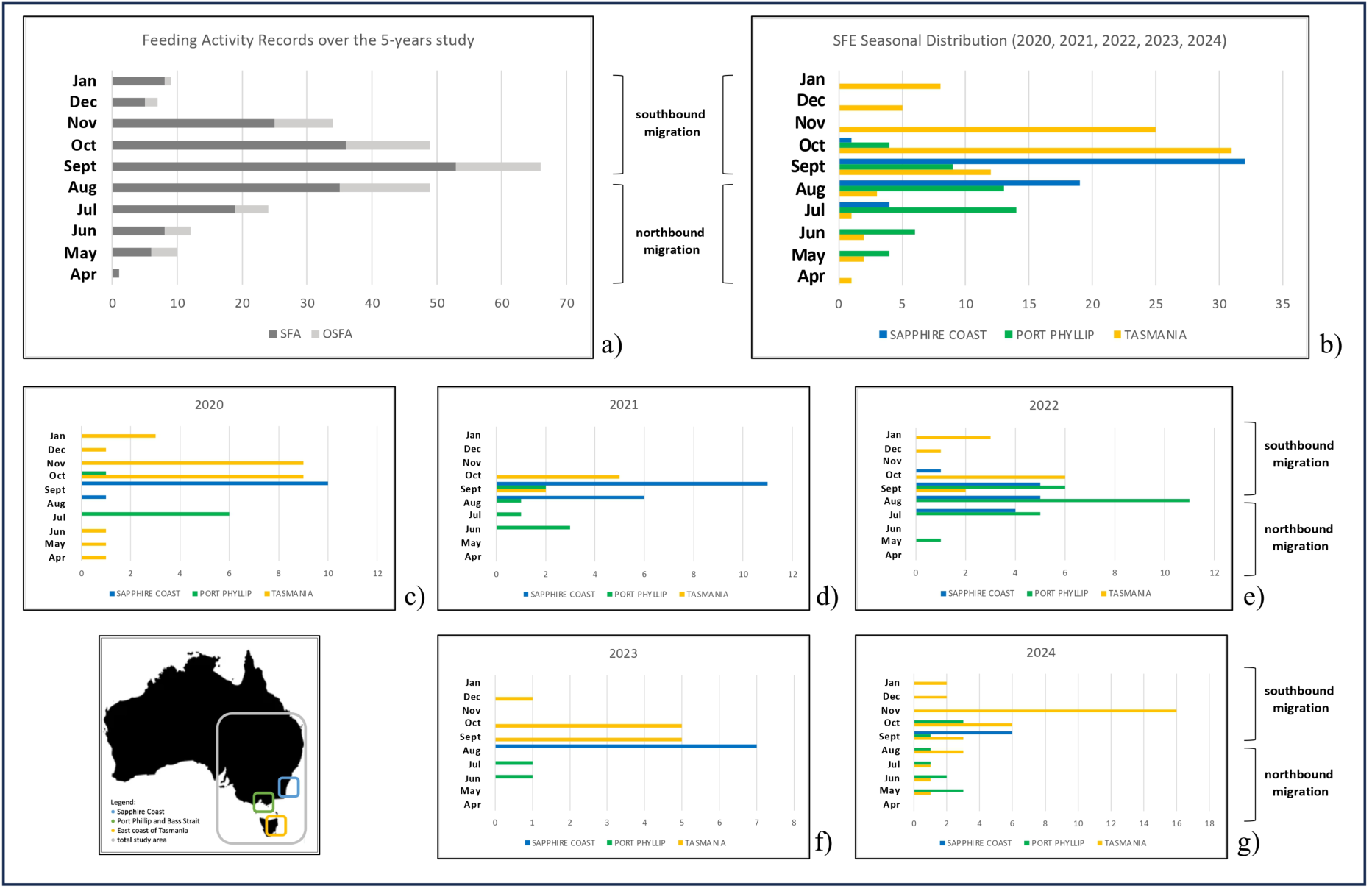
Bar charts of humpback whale feeding events records across the five study years (2020–2024), categorized by migratory months and geographic regions: a) total Supplementary Feeding Events (SFE) counts (n=256), both within proposed Supplementary Feeding Areas (SFA) and Outside proposed Supplementary Feeding Areas (OSFA) plotted per migratory month; b) Seasonal distribution of SFE (2020–2024) in the three proposed Supplementary Feeding Areas (SFA): Sapphire Coast, Port Phillip and Bass Strait, and East Coast of Tasmania (n=193); c), d), e), f) and g) SFE counts, broken down by region and migratory month for years 2020 (n=40), 2021 (n=34), 2022 (n=47), 2023 (n=20) and 2024 (n=52) respectively. An inset map of Australia identifies the study region and the three proposed Supplementary Feeding Areas (SFAs).

### Data quality control

A *post hoc* quality-control audit was conducted to identify potential duplicate records. Initial screening identified nine candidate duplicate pairs sharing the same observation date and reported locality. Each pair was subsequently examined using all available contextual information. The candidate pairs differed substantially in reported pod size and/or lacked sufficient metadata (e.g., observation time or precise location) to determine whether they represented repeated reports of the same feeding event or independent feeding events occurring within the same locality on the same day. Consequently, no records were merged or excluded from the final dataset.

### Pod size during feeding activities

Feeding records in the proposed SFAs were categorized with regard to the number of whales involved in the feeding activity. Feeding in a pod (defined as at least two humpback whales within five meters of each other) was commoner than feeding as a singleton in the years 2020, 2021, and 2023, while in 2022 and 2024, the singletons and feeding pods occurred in roughly equal proportions (Table 1 and Figure 3a).

**Figure 3.**
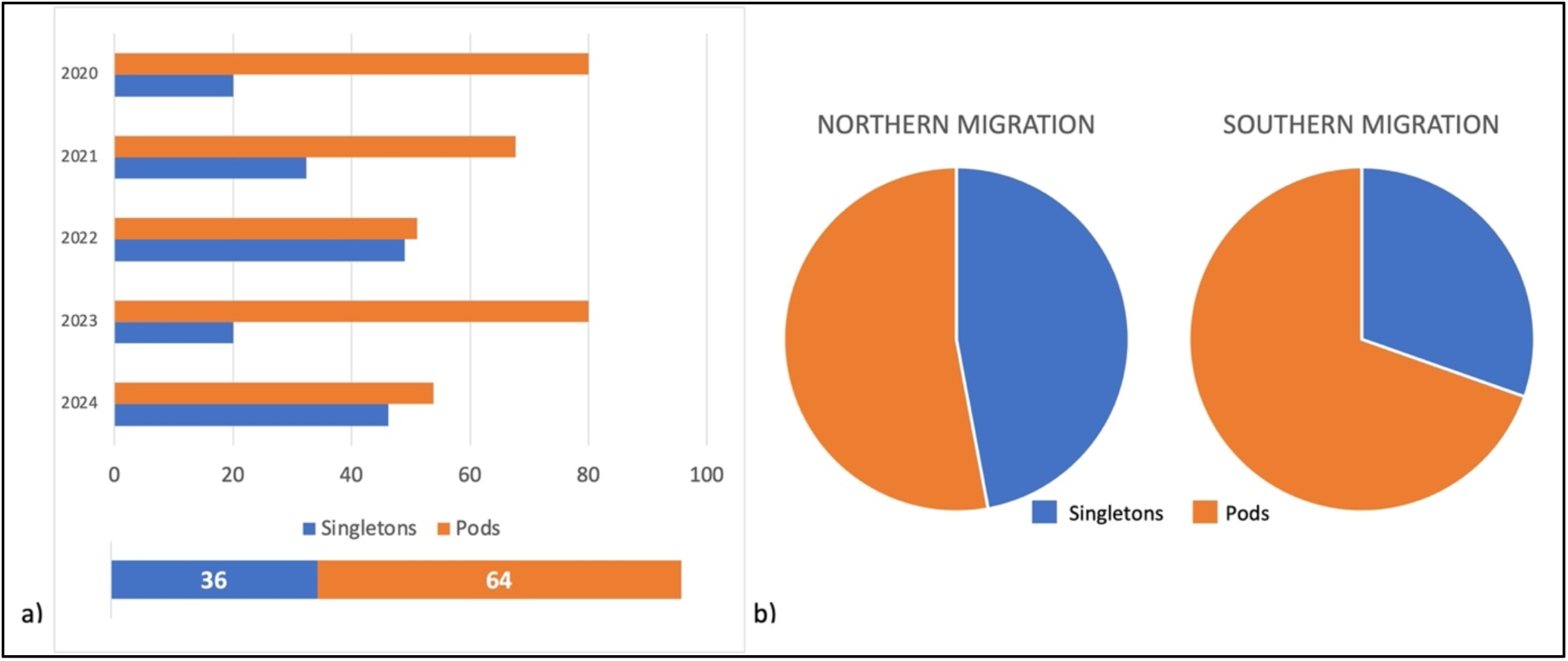
Percentage of singletons’ and groups’ SFE recorded between 2020 and 2024 (a). In the figure b. they are divided between the Northern Migration (considered as the humpback whales migrating from April till August) and the southern migration (considered as the humpback whales migrating from September till the first half of January).

**Table 1.** Singletons vs groups of humpback whales sighted feeding in Eastern Australia over the study period. All sightings were split into those where just one single humpback whale was spotted feeding (1) and the ones where a pod of humpback whales was seen feeding (1+). The sightings were divided between months on the columns and years on the rows. The final column presents the total number of sightings divided per year, while the bottom row reports the total number of sightings divided per month.

|  | April |  | May |  | June |  | July |  | August |  | September |  | October |  | November |  | December |  | January |  | TOT |  |
| --- | --- | --- | --- | --- | --- | --- | --- | --- | --- | --- | --- | --- | --- | --- | --- | --- | --- | --- | --- | --- | --- | --- |
| Yearly migration | 1 | 1+ | 1 | 1+ | 1 | 1+ | 1 | 1+ | 1 | 1+ | 1 | 1+ | 1 | 1+ | 1 | 1+ | 1 | 1+ | 1 | 1+ | 1 | 1+ |
| 2020 | 1 | 0 | 1 | 0 | 0 | 1 | 3 | 3 | 0 | 1 | 1 | 9 | 1 | 9 | 0 | 9 | 1 | 0 | 0 | 0 | 8 | 32 |
| 2021 | 0 | 0 | 0 | 0 | 2 | 1 | 1 | 0 | 4 | 2 | 2 | 14 | 1 | 4 | 0 | 0 | 0 | 0 | 1 | 2 | 11 | 23 |
| 2022 | 0 | 0 | 1 | 0 | 0 | 0 | 3 | 6 | 8 | 8 | 7 | 6 | 3 | 4 | 0 | 0 | 1 | 0 | 0 | 0 | 23 | 24 |
| 2023 | 0 | 0 | 0 | 0 | 0 | 1 | 0 | 1 | 0 | 7 | 1 | 4 | 2 | 3 | 0 | 0 | 1 | 0 | 0 | 0 | 4 | 16 |
| 2024 | 0 | 0 | 3 | 1 | 1 | 2 | 0 | 2 | 4 | 0 | 3 | 7 | 4 | 5 | 5 | 11 | 2 | 0 | 2 | 0 | 24 | 28 |
| TOT | 1 | 0 | 5 | 1 | 3 | 5 | 7 | 12 | 16 | 18 | 14 | 40 | 11 | 25 | 5 | 20 | 5 | 0 | 3 | 2 | 70 | 123 |

However, when data from the five years were pooled together, the observed frequencies of singleton and pod feeding events during the northern and southern migrations (Table 2) were compared using a Chi-square test. The analysis yielded a significant difference between migration phases (χ² = 5.29, d.f. = 1, p = 0.0215). All expected cell frequencies exceeded 5, confirming that the assumptions of the test were met. The chi-square test was performed without Yates’ continuity correction.

**Table 2.**
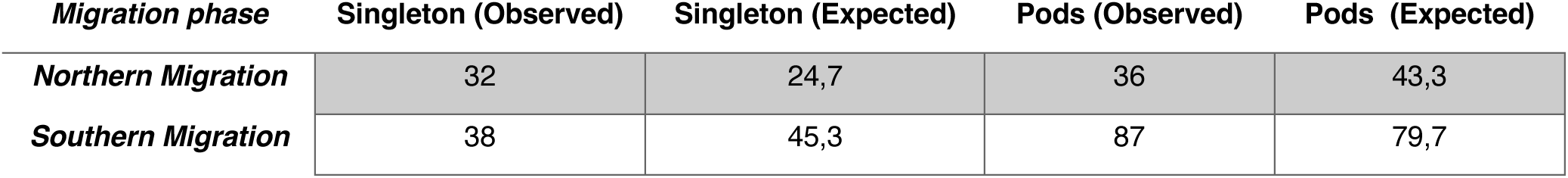
2 × 2 contingency table. Observed and expected frequencies used for the chi-square analysis.

Since the p-value is less than the significance level of 0.05, we can reject the null hypothesis and conclude that there are significant differences in pod size of feeding whales during the two migrations, namely encountering considerably more feeding whale aggregations during the southern migration (Figure 3b).

For 98 out of the 193 feeding events (50.77%), it was possible to observe the feeding behavior of humpback whales using the Eastern Australia migratory corridor during the five analyzed years (Figure 4), and for 54 out of 193 feeding events (27.97%), also the type of prey consumed (Figure 5) during candidate supplementary feeding activities. The results are based on the sightings where it was possible to define the feeding strategies (lunge or bubble net) employed or the type of prey item (fish or krill) ingested.

**Figure 4.**
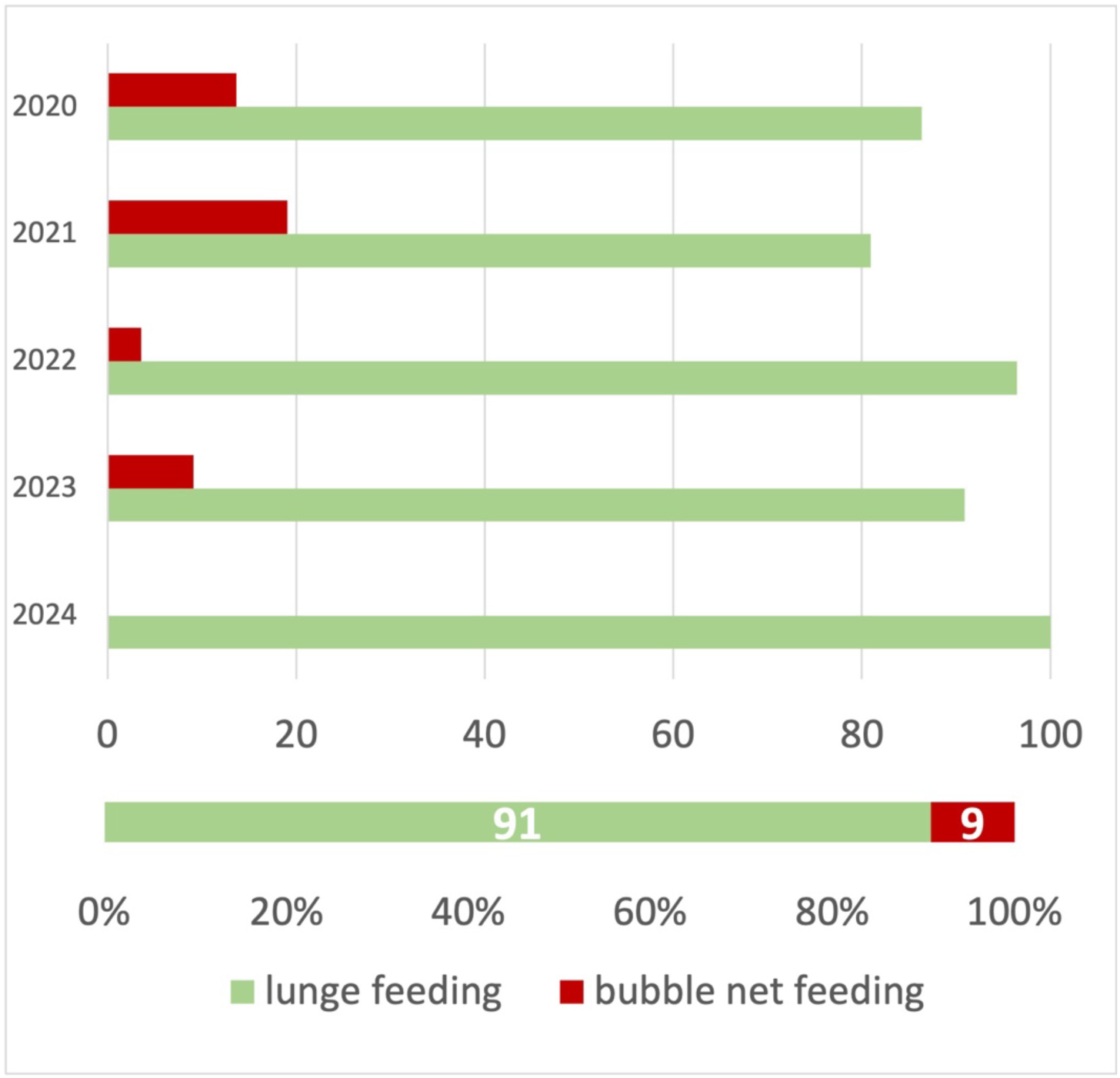
“Lunge feeding” and “Bubble net feeding” percentages recorded during humpback whales SFE in each of the five years (2020-2024) and in the overall sample set.

**Figure 5.**
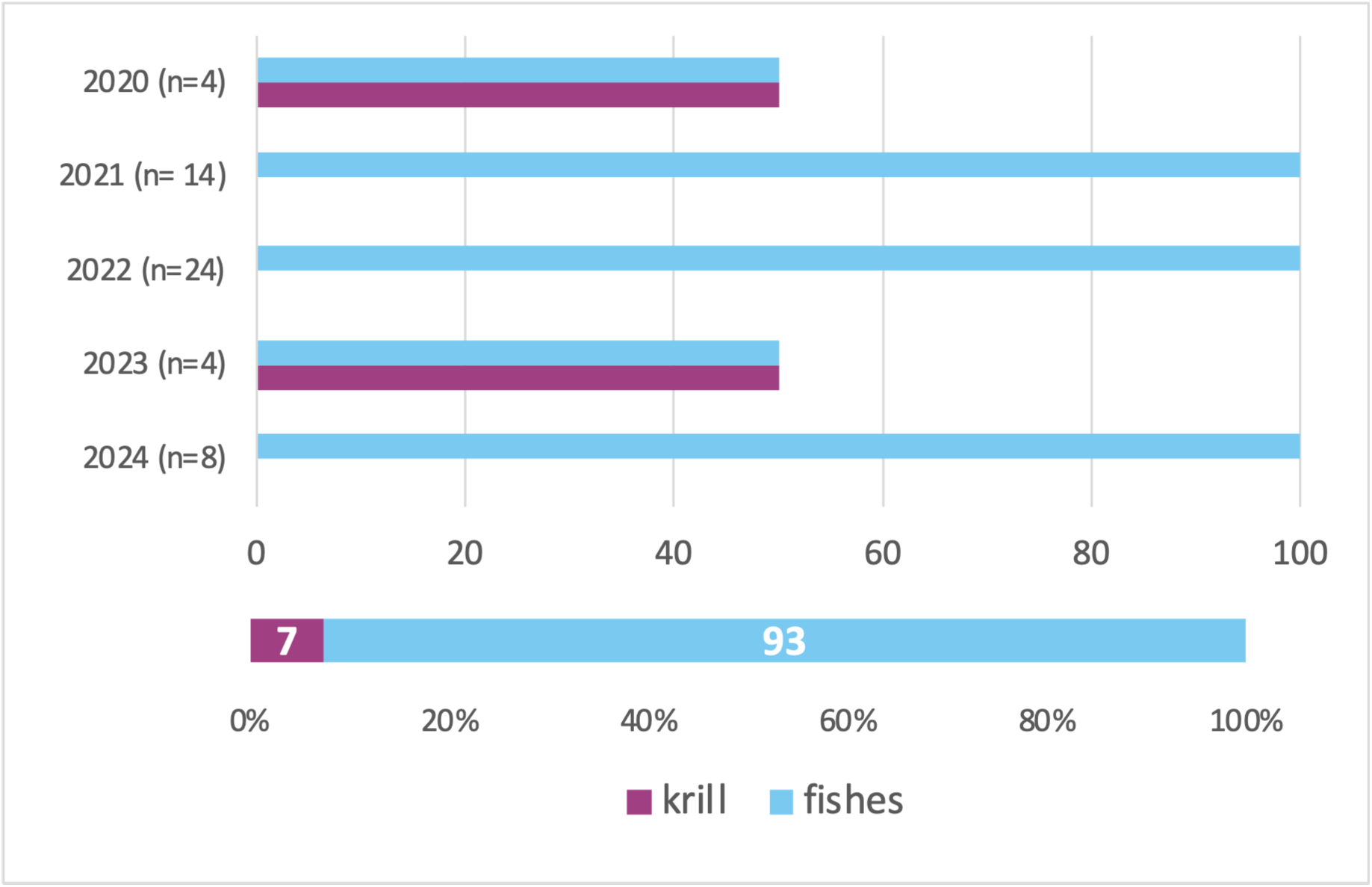
Proportion of humpback whale SFE categorized by prey type (krill vs. schooling fishes) shown by year (2020 to 2024) and in the overall sample.

### Comparison between passive and active citizen science

Of the 256 feeding records collected, 196 (76.6%) were retrieved through passive citizen science using Facebook and Instagram. For the purpose of this comparison, five records originally published as online news articles but retrieved through these social media platforms were reported separately from the remaining 191 user-generated social media records. The remaining 60 records (23.4%) were obtained from the active citizen science platform Happywhale (Table 3). Feeding behaviour could be identified in 118 of the 191 social-media records (61.8%) compared with 17 of the 60 Happywhale records (28.3%). In contrast, prey type could be identified in 53 social-media records (27.7%) and 15 Happywhale records (25.0%), with no significant difference between the two datasets.

**Table 3.**
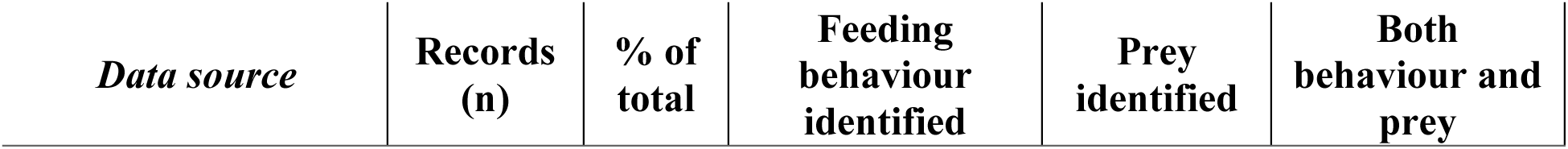
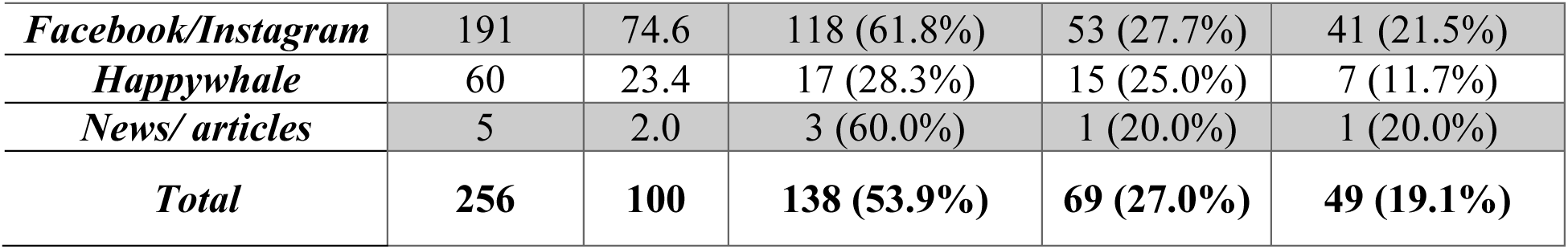
Comparison of information obtained from passive and active citizen science sources. For this comparison only, five records originating from online news articles but retrieved through Facebook or Instagram were treated as a separate category to distinguish journalistic content from user-generated social media posts. In all other analyses, these records were included within the Facebook/Instagram dataset because they were retrieved through these social media platforms.

**Table 4.**
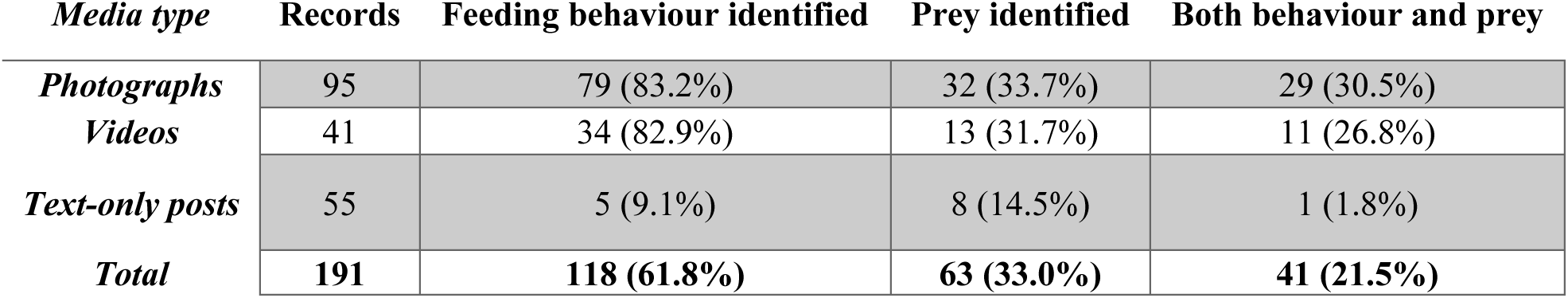
Information obtained according to media type (Facebook and Instagram records only)

### Media type

Among the 191 records obtained from Facebook and Instagram, 95 consisted of photographs, 41 of videos and 55 of text-only posts (Table Y). Feeding behaviour was identified in 79 photographic records (83.2%), 34 video records (82.9%) and only five text-only posts (9.1%). The proportion of records allowing behavioural identification differed significantly among media types. Conversely, prey type was identified in 32 photographic records (33.7%), 13 video records (31.7%) and eight text-only posts (14.5%), with no significant differences among media types.

## DISCUSSION

### Passive Versus Active Citizen Science

This study explores the potential of opportunistic crowdsourced data from social media as a tool to investigate overlooked aspects of whale ecology, such as feeding behaviour, and compares this “passive” citizen science (CS) approach with the more widespread “active” one using the same data source (social-media photographs) retrieved through one of the most established whale-focused CS platforms, the Happywhale network.

The comparison between passive and active citizen science datasets highlights an important difference in the type of ecological information that can be extracted from each source. Although Happywhale contributed valuable occurrence records, feeding behaviour was identified significantly more frequently in records obtained from Facebook and Instagram than in those retrieved from Happywhale. This likely reflects the different primary purposes of the two platforms. Happywhale is specifically designed for photo-identification, where users are encouraged to upload images of the ventral surface of the tail flukes for individual recognition. Consequently, behavioural context is often absent or limited. In contrast, social media users frequently share photographs or videos documenting unusual or engaging events, often accompanied by descriptive captions, thereby preserving ecological and behavioural information that might otherwise be lost.

Prey identification, however, showed a different pattern from feeding behaviour detection. Interestingly, the proportion of records allowing prey identification did not differ significantly between passive social-media records and Happywhale submissions. This suggests that while passive citizen science provides a clear advantage for documenting feeding behaviour, the identification of prey remains constrained by factors such as image quality, viewing angle and prey visibility, regardless of the platform from which the records originate. These constraints were also evident when comparing the different media types within the passive citizen science dataset.

Within the passive citizen science dataset, photographs and videos were similarly effective for documenting feeding behaviour, whereas text-only posts rarely contained sufficient information for behavioural classification. This finding suggests that a single well-timed photograph may often provide enough visual evidence to identify conspicuous feeding techniques, despite the greater temporal information generally available from videos. Conversely, prey identification remained comparatively limited across all media types, reflecting the inherent difficulty of recognising prey species during feeding events. Taken together, these results highlight the broader value of passive citizen science for behavioural ecology.

Overall, these findings support the value of passive citizen science as a complementary source of behavioural information.Passive citizen science contributes not only additional occurrence records, but also behavioural context that is frequently absent from structured citizen-science platforms. The combination of both approaches therefore provides a more complete description of humpback whale feeding ecology than either source alone. This complementary value provides the basis for interpreting the ecological patterns identified in the present study.

### Identification of Potential Supplementary Feeding Areas

This study demonstrates that sporadic, ‘opportunistic’ photographic material can sometimes yield more informative insights into underexplored aspects of whale ecology than systematically collected image sets intended primarily for photo-identification (Dickinson et al. 2010; Di Minin et al. 2015; Jarić et al. 2020; Toivonen et al. 2019). In the case of the humpback whale, such targeted collections typically focus on photographing the ventral surface of the tail fluke. Consequently, other images that may capture different aspects of the animal’s life are often unanalysed (Franklin et al. 2020; Cheeseman et al. 2023).

Although the dataset is relatively limited, it nevertheless provides a useful proof of concept. The study focused on data from a limited area (Eastern Australian coast), over a relatively short time span -five migratory seasons- and uses as a source only two of the most popular social media (Facebook and Instagram). Yet, the present work shows how this type of spontaneous data can be useful for getting preliminary information on unexplored wildlife aspects, taking advantage of the increasing willingness of people to voluntarily share information and photographs online. In our specific case, data retrieval was undoubtedly facilitated by the fact that whales in general are charismatic creatures and that the numerous whale-watching programs spread globally create an easy opportunity for the general public to meet these fascinating giants of the seas. This proof of concept is reflected in the identification of recurrent feeding areas along the migratory corridor.

Indeed, this exploratory study, subject of a master’s thesis project, was nevertheless able to draw attention to three specific and well-defined areas of interest along the migratory route of E1 humpback whales which are the Sapphire Coast, Port Phillip/Bass Strait, and South-eastern Tasmania, likely to represent elective supplementary feeding areas (SFAs), all three located in the southernmost portion of the Eastern Australian migratory corridor. Interestingly, although sporadic feeding activity was reported as north as Hervey Bay (around −24.89 degrees of latitude), feeding events fulfilling the “supplementary feeding” criteria were recorded predominantly in the southernmost portion of the Eastern Australia migratory corridor (southern than −32.77 degrees of latitude). Moreover, candidate supplementary feeding events were also twice as abundant during the southern (n=125, 64.77%) rather than the northern (n=68, 35.23%) migration.

Results are not surprising if we consider that, for instance, Tasmania is a renowned high-productivity area (e.g., Kämpf 2015). Moreover, these areas, especially the Sapphire Coast and Tasmania, have already come to the attention of scientists because of previous sightings of occasional feeding humpback whales (Gill et al. 1998; Stamation et al. 2007; Owen et al. 2015; Owen et al. 2017; Pirotta et al. 2021). Thus, our results support previous observations, while also suggesting that putative supplementary feeding areas may be more extensive than indicated by the current dataset.

### Feeding strategies and social context of supplementary feeding

Although not originally collected for behavioural analyses, these opportunistic observations provided an opportunity to explore feeding strategies and social patterns associated with putative supplementary feeding events.

In the migratory seasons of 2020, 2021, and 2023, there was a clear predominance of pods (considered as groups composed of more than one humpback whale) engaged in feeding activity rather than single individuals. This observation does not necessarily reflect a tendency towards gregarious foraging but more likely reflects the typical social composition of humpback whales during migration and the breeding season, when individuals interact socially for reproductive purposes and groups therefore predominate over singletons (e.g., Valsecchi et al. 2002).

Interestingly, feeding pods were particularly prevalent during the southern migration. This pattern may reflect the naturally higher occurrence of social groups during this migratory phase, including mother– calf associations (McGovern et al. 2025). Alternatively, multiple individuals may temporarily aggregate while exploiting the same feeding areas. The lower occurrence of feeding groups in May, December, and January may instead reflect reduced migratory intensity during these periods, suggesting that humpback whales exploit feeding opportunities regardless of whether they migrate alone or in association with conspecifics.

At the same time, the social interpretation of these observations should be treated with caution. The predominance of pods during southern migration may also indicate that multiple individuals respond simultaneously to local feeding opportunities. However, because observations were derived from passive citizen science and group composition could not always be verified, temporary spatial overlap of independently migrating whales cannot be excluded.

Lunge feeding behaviour (including both lateral and vertical lunge feeding) was the predominant feeding strategy observed and is consistent with what is already known for E1 humpback whales in traditional feeding areas (Stamation et al. 2007; Owen et al. 2015; Owen et al. 2016; Owen et al. 2017).

Prey items included both krill and fish with almost equal frequency overall; however, in 2021, 2022, and 2024, humpback whales appeared to forage exclusively on fish within Australian supplementary feeding areas. One possible explanation is that reduced krill availability promoted opportunistic feeding on locally abundant fish schools. This interpretation is consistent with the migratory timing of E1 humpback whales, which reach Australian waters during the Southern Hemisphere winter, when krill availability begins to decline following the autumn peak in density and biomass (Young et al. 1993).

Analyzing the total number of feeding humpback whales across the five study years, megapods were more frequently observed in 2020 and 2021 than in subsequent years. This pattern may reflect the unusual conditions associated with the COVID-19 pandemic, during which reduced maritime traffic and lower fishing activity may have resulted in quieter waters and altered local ecological conditions. However, changes in observer activity and social-media posting behaviour during COVID-19 restrictions may also have contributed to the apparent increase in megapod records. Therefore, this interpretation remains speculative and would require dedicated investigation.

### Environmental drivers of supplementary feeding

Over the past few years, climate change started degrading Antarctic feeding areas, leading to lower nutrient concentrations (Loeb and Santora 2015; Atkinson et al. 2019) and, consequently, lower energy intake by humpback whales before they start their migration to the breeding grounds. Reduced energetic intake prior to migration may increase energetic constraints during migration and could contribute to the occurrence of supplementary feeding behaviours. Interestingly, the majority of the feeding events reported in this study are recorded during the southbound migration, when energy reserves may become increasingly depleted, and at latitudes ranges (36.2° S - 43.4° S) intermediate between feeding and breeding grounds. A few scientists have also hypothesized that due to the climatic changes, there could be a gradual switch from the Antarctic feeding areas to other feeding areas (Loeb et al. 1997; Atkinson et al. 2004; Flores et al. 2012; Kawaguchi et al. 2013; Eisenmann et al. 2016). To date, this switch has not been observed in Australia.

However, different studies have highlighted how climate change and La Niña have been affecting Australia’s climate since September 2020 (Supplementary Figures 1 and 2), including an increase in the average rainfall (Supplementary Figure 2) resulting in greater river discharges and an increase of nutrients in the oceans (Skerratt et al. 2019) which could explain the increasing number of feeding events spotted.

It is not surprising that the incidence of target records in each of the three supposed supplementary feeding areas varies significantly across the five years analysed, with, for example, Tasmania scoring a greater number of feeding sightings in 2020, probably reflecting the documented strong interannual changes in the timing of the nitrate decline, and thus of phytoplankton spring bloom, whose timing can vary from year to year by as much as four months (Harris et al. 1987)

Overall, these patterns suggest that environmental variability, including climate-driven changes and interannual oceanographic fluctuations, may play an important role in shaping the occurrence and distribution of possible supplementary feeding events along the Eastern Australian migratory corridor. However, causality cannot be established from the present dataset, and continued long-term monitoring will be essential to determine whether these patterns become increasingly frequent under future environmental change.

### Study advantages and limitations

Unlike conventional whale surveys, passive citizen science exploits observations that already exist and therefore represents an essentially cost-free and widely accessible monitoring infrastructure. The approach demonstrated here highlights the potential of social-media-derived observations to provide broad-scale ecological information relevant to marine mammal conservation and management.

Passive citizen science offers several practical advantages for marine mammal monitoring. Because data originate from observations spontaneously collected and shared by members of the public, this approach requires no dedicated field campaigns or specialised equipment and can provide extensive spatial and temporal coverage. In addition, rare or unexpected behaviours may be detected due to the large number of potential observers distributed across broad geographic areas. In the specific context of data opportunistically collected by whale-watching customers, this proof of concept demonstrates that their participation can yield valuable scientific information. This potential could be leveraged by whale-watching companies, which might proactively encourage their customers to share images of their experiences while providing as much contextual information as possible (including date, time, and location). This simple yet effective measure would greatly enhance the amount of usable data extractable from such publicly shared materials.

Similar advantages of opportunistic citizen science have been recognised across a wide range of ecological applications, particularly for documenting rare events, expanding geographic coverage and generating novel ecological hypotheses (Dickinson et al. 2010; Bonney et al. 2014; Chandler et al. 2017). Future developments could further enhance its effectiveness through automated image and video analysis using machine learning and artificial intelligence, potentially improving the scalability and efficiency of data processing (Weinstein 2018; Christin et al. 2019). In a broader context, citizen-derived observations could be integrated with complementary ecological datasets, including remotely sensed oceanographic variables, prey distribution and habitat models, could provide an increasingly powerful framework for investigating marine predator ecology at large spatial scales (Toivonen et al. 2019; Jarić et al. 2020). Taken together, these developments highlight the increasing potential of passive citizen science as part of a wider multi-platform approach to marine biodiversity monitoring.

Nevertheless, passive citizen science is also associated with important limitations. Unaware participants were not trained to collect standardized, relevant and complete information for this research. This implies that the data could be incomplete and inconsistent across participants. Moreover, behavioural interpretations may be constrained by the quality of photographs or videos available and social-media accessibility may introduce spatial and demographic biases potentially leading to uneven sampling effort across regions and years (Dickinson et al. 2010; Kosmala et al. 2016; Isaac et al. 2014). To further contextualise this potential bias, whale-watching operator distribution was compared among regions as a qualitative proxy of observation opportunity (Supplementary Table S2). This comparison indicated that the regions with the highest number of feeding records did not correspond to those with the highest number of whale-watching operators (e.g. Harvey Bay or Gold Coast), suggesting that the spatial patterns identified in this study are unlikely to be explained solely by differences in observation opportunity. Nevertheless, passive citizen science should be viewed as a complementary monitoring tool rather than a replacement for systematic survey programmes.

Although each feeding event was treated as an independent observation for the statistical analyses, we acknowledge that some events recorded on consecutive days may have involved the same whale groups, particularly during prolonged feeding aggregations such as the Bermagui megapods. Consequently, complete independence among observations cannot be guaranteed, and this potential pseudo-replication should be considered when interpreting the chi-square results.

Another limitation concerns the possibility of residual duplicate records, an inherent challenge when integrating passive citizen-science observations originating from multiple independent contributors and social-media platforms. To minimise this source of bias, all potentially overlapping records were manually evaluated during dataset compilation using observation date, reported locality, pod size, observation time (when available) and all additional contextual information. A subsequent *post hoc* quality-control audit identified only nine candidate duplicate pairs among the 256 records (3.5% of the dataset). Of these candidate pairs, all were retained because the available metadata were insufficient to distinguish repeated reports of the same feeding event from independent feeding events occurring within the same reported locality on the same day. Moreover, reported localities generally referred to broad coastal areas rather than precise observation sites, allowing multiple independent feeding events to legitimately occur within the same locality within a single day. Consequently, any residual duplication is expected to introduce only a slight positive bias in the total number of feeding events and is unlikely to substantially influence the identification of the proposed Supplementary Feeding Areas.

Despite these limitations, the present study demonstrates how passive citizen science can contribute to identifying emerging ecological phenomena and generating hypotheses for future targeted monitoring. Rather than replacing systematic survey programmes, passive citizen science should be viewed as a complementary monitoring approach capable of revealing ecological patterns that may otherwise remain undetected (Bonney et al. 2014; Chandler et al. 2017; Jarić et al. 2020). In the present study, the recurrent spatial patterns identified in the present study suggest that passive citizen science may provide a useful early-detection tool for identifying emerging feeding areas and behavioural shifts in migratory humpback whales.

## CONCLUSIONS

This study provides a pilot demonstration of the potential of passive citizen science as an innovative and complementary methodological approach for the collection of ecologically relevant observations, highlighting its capacity to generate preliminary scientific hypotheses from opportunistic, publicly available data. Compared to active citizen science frameworks, this approach can reduce data collection constraints linked to predefined sampling designs, while still providing valuable broad-scale ecological information for marine mammal research.

The presented data suggest the presence of three possible supplementary feeding areas (SFAs) for the E1 population of humpback whales, which correspond to highly productive and nutrient-rich zones on the Sapphire Coast (New South Wales), in southern Australia (off Melbourne from Point Lonsdale to Wilsons Promontory), and on the southeast coast of Tasmania. The identification of these areas of recurrent feeding suggests that supplementary feeding activity may be more widespread than currently documented and potentially underestimated, particularly where opportunistic feeding events (OFE) have been recorded but not yet deeply analysed due to limited sampling effort. Given the methodological nature of this study, the results also highlight the importance of improving spatial and temporal coverage of observations to refine the boundaries and ecological relevance of these candidate feeding areas. In this context, integrating passive citizen science outputs with targeted field approaches, such as boat-based surveys and individual identification through Photo ID (for example integrating citizen science records from the Happywhale database), could help assess site fidelity and potential inter-annual return patterns of individuals to specific feeding regions, as well as the demographic structure of feeding whales, including, where possible, demographic structure (e.g., age class and, when inferable, sex), particularly in relation to mother–calf associations, which are particularly affected by energetic constraints during migration (McGovern et al. 2025).

This study simplifies complex ecological patterns to highlight the occurrence of putative supplementary feeding events, demonstrating how passive citizen science can effectively contribute to the detection of emerging behaviours and the formulation of testable hypotheses. Overall, passive citizen science based on images and annotations shared by media users from all over the world can represent a continuously expanding and scalable methodological tool, capable of supporting large-scale, continuous monitoring of underexplored ecological phenomena and contributing to early detection of behavioural changes in marine ecosystems.

## Supporting information

Supplementary images and Supplementary Tables

## ACKNOWLEDGEMENTS

We would like to acknowledge Griffith University, and in particular Sarah McCullock and Olaf Meynecke, for the initial discussions and general suggestions that helped identify the research topic subsequently explored in the author’s Master’s thesis. The Master’s thesis provided an initial basis for the present research, which the authors subsequently further developed and expanded.

We also thank the members of the public who shared wildlife observations and photographs through social media, as well as the contributors to Happywhale, whose publicly available records provided valuable information for this study.

