## Supplementary images and Supplementary Tables for "World Wide Eyes: Social Media Assisted Wildlife Research. Assessing Supplementary Feeding Areas for Eastern Australian Humpback Whales as a Case Study"

**G. Marzorati^1,^ & E. Valsecchi^1,2*^**

^1^ Department of Environmental and Earth Sciences, University of Milano-Bicocca, Piazza della Scienza 1, 20126, Milan, Italy

^2^ MaRHE Center, Magoodhoo Island, Faafu Atoll, Republic of Maldives

**KEYWORDS:**  *Megaptera novaeangliae,* passive citizen science, social media platforms,

participative data, wildlife sightings, nature photography

**SUPPLEMENTARY MATERIAL**

**Supplementary Figure 1.** *Time Series of the ENSO index from NOAA for 1950 to Feb 2025. The index is calculated using the sea surface temperatures in the Niño 3.4 region (area between 170°W-120°W and 5°S-5°N). The dashed red line shows the temperature threshold (NOAA uses +0.5 Celsius), and El Niño events are shaded in red. The strongest events occurred in 1997/98 and 2015/16 and are shaded in blue. The 2002/03 weak El Niño, shaded in blue, caused widespread impacts on Australia. Source: ARC Centre of Excellence for Climate Extremes.*

**Supplementary Figure 2.** *Yearly historical records of Australia's rainfall (top) and temperatures (bottom). Source: images retrieved from http://www.bom.gov.au/climate/history/rainfall/ and httip://www.bom.gov.au/climate/history/temperature/,* respectively*.*

**Supplementary Table 1.** List of humpback whale feeding records in East Australia between 2020 and

2025.

**Supplementary Table 2.** **Compilation of commercial whale-watching operators used as a qualitative proxy of observation opportunity.** To account for potential spatial variation in public observation opportunity, we compiled the documented distribution of commercial whale-watching operators active along the eastern and southern Australian coastline during the study period (2020–2025). Operators were identified through publicly available sources, including official operator websites, regional tourism information portals, and governmental or conservation-related resources where available. Operators were included if they (i) explicitly advertised dedicated whale-watching activities, (ii) were active at any point during the study period, and (iii) operated within the geographical area considered in this study.

Each commercial operator was counted once, independently of the number of vessels, departure ports, or tour products offered, to avoid artificially inflating observation opportunity in regions where individual companies operated multiple vessels or locations. When operators served multiple regions, they were assigned to all relevant regions in the dataset. In cases of company name changes, acquisitions, or changes in ownership during the study period, operators were retained separately when they represented distinct periods of commercial activity. The resulting operator distribution was used as a qualitative indicator of relative observation opportunity and does not represent a direct measure of sampling effort, observation hours, vessel activity, or total whale-watching intensity.

**Supplementary Table 3.** **Regional distribution of humpback whale feeding records, whale-watching operators, and qualitative observation effort proxy.** The table compares the number of feeding records collected in each study region with the number of documented commercial whale-watching operators identified in Supplementary Table S1. Whale-watching operator distribution was used as a qualitative proxy to contextualise potential spatial differences in observation opportunity associated with passive citizen science data.

Observation effort categories were assigned according to the number of documented whale-watching operators operating within each region: Low (1–2 operators), Moderate (3–4 operators), High (5–6 operators), and Very High (≥7 operators). This classification was used only to evaluate potential spatial variation in observation opportunity and was not applied as a correction factor or normalization of feeding records. Because passive citizen science observations derive from multiple sources, including whale-watchers, recreational observers, and independent contributors, operator numbers should be interpreted as a relative proxy rather than a complete estimate of total observation effort.

Supplementary Figure 1.

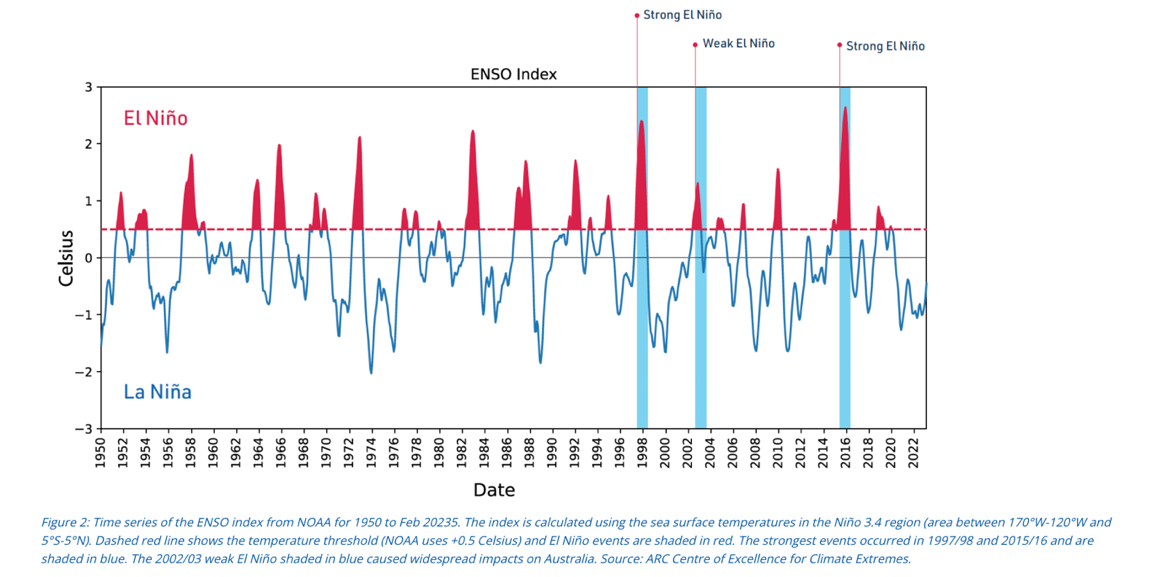

Supplementary Figure 2.

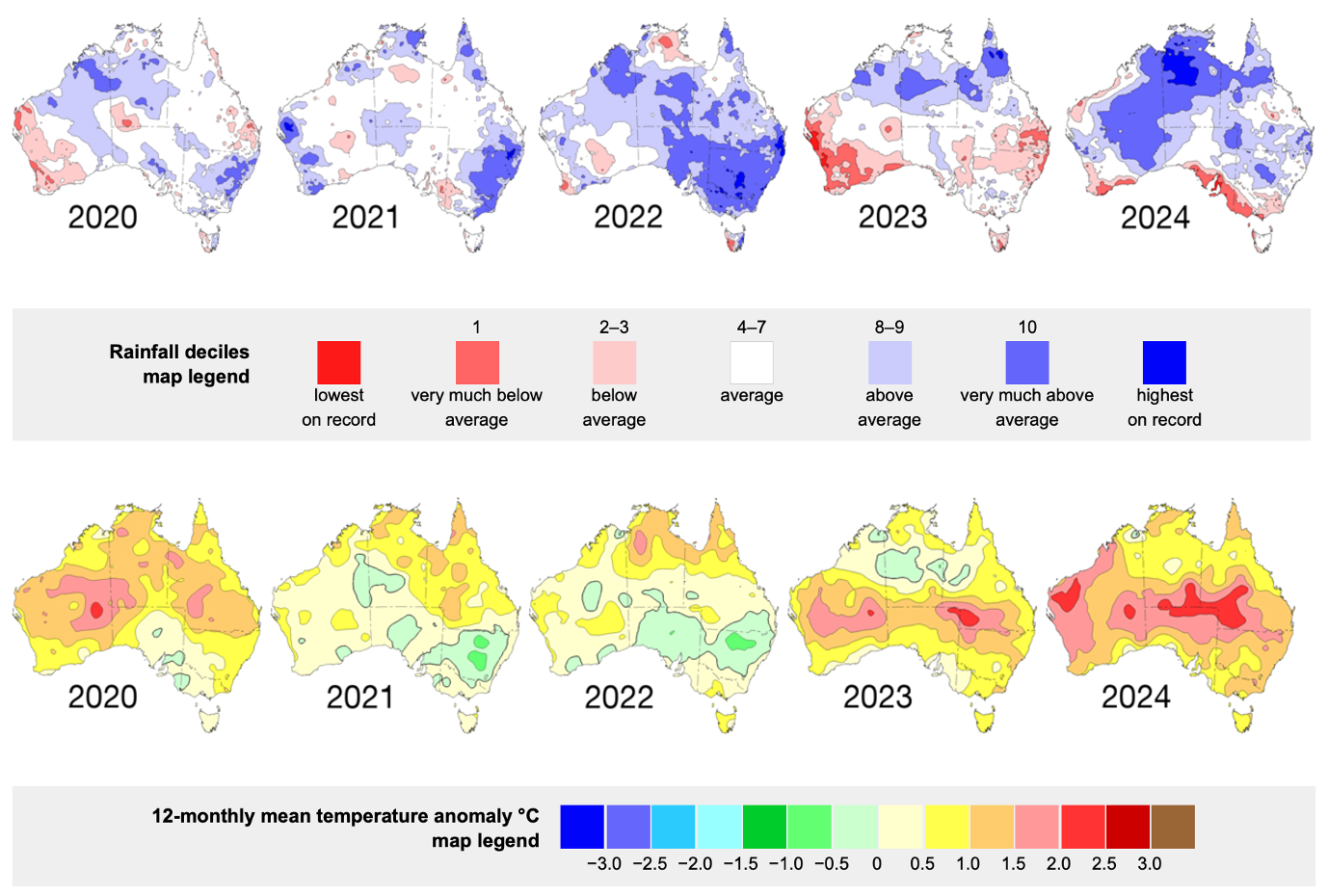

TABLE S1.

| Date | Location | Coordinates | | | Time | N° of whales | Type of evidence | Source | Type of feeding | type of food |
| --- | --- | --- | --- | --- | --- | --- | --- | --- | --- | --- |
| 23/04/20 | Tasman Peninsula | -42.934884  147.746565 | | |  | 1 | post | Facebook | \ | \ |
| 17/05/20 | Cape Pillar | -43.190253  147.994761 | | |  | 1 | post | facebook | \ | pyrosomes and krill |
| 06/06/20 | Prossers Bay | -42.556448  147.895126 | | | evening | 1+ | post | facebook | \ |  |
| 7/07/2020 | Wilsons Promontory | -39.142472  146.554293 | | | afternoon | 8-…-10 | photo | facebook | \ |  |
| 07/07/20 | Pyramid Rock (Phillip Island) | -38.554639  145.231893 | | |  | 1 | photoes | Facebook | Lunge feeding | \ |
| 08/07/20 | Phillip Island | -38.566168  145.381362 | | |  | 2 | photo | Facebook | vertical lunge feeding | \ |
| 08/07/20 | Pedra Branca | -43.850126  146.969027 | | |  | 1+ | post | facebook | \ | tuna |
| 9/07/2020 | Phillip Island | -38.101772  144.898041 | | |  | 1 | post | facebook | Vertical lunge  feeding |  |
| 9/07/2020 | Pyramid Rock | -38.530112  145.226568 | | |  | 1 | photoes | facebook | lunge feeding | Baitfish |
| 26/07/2020 | Pyramid Rock (3 km south) | -38.611398  145.224216 | | | 10:15 AM | 3 | video | facebook | \ |  |
| 03/08/20 | Sydney (Long Reef) | -33.74701919  151.30528180 | | | Afternoon | 1 | photoes | Facebook | Lunge feeding | \ |
| 03/08/20 | Sydney | -33.74701919  151.30528180 | | | Afternoon | 2 | photoes | Facebook | Lunge feeding | \ |
| 04/08/20 | Sydney | -33.874472  151.330916 | | |  | 1 | video | Facebook | Lunge feeding | \ |
| 07/08/20 | Bass Point | -34.598418  150.906184 | | |  | 2 | photoes | Facebook | \ | \ |
| 09/08/20 | Sydney (Long Reef) | -33.74701919  151.30528180 | | | Sunset | 1 | photoes | Facebook | Lunge feeding | \ |
| 30/08/20 | Eden (NSW) | -37.069081  149.941270 | | |  | 2 | post | Facebook | Lunge feeding | \ |
| 05/09/20 | Eden (NSW) | -37.061687  149.969912 | | |  | 1+ | \ | Facebook | \ | \ |
| 06/09/20 | Eden (New South Wales) | -37.043697  149.977697 | | |  | 100 | **photoes** | instagram | lunge feeding |  |
| 11/09/20 | Tamarama (NSW) | -33.900403  151.273057 | | |  | 1 | post | Facebook | Lunge feeding | \ |
| 12/09/20 | Merimbula (New South Wales) | -36.913467  149.927622 | | | 9:30-12:30 | 100+ | \ | instagram | lunge feeding |  |
| 12/09/20 | Eden (New South Wales) | -37.085612  149.912809 | | |  | 2+ | photoes | instagram | vertical lunge feeding |  |
| 12/09/20 | Eden (New South Wales) | -37.075752  149.955724 | | |  | megapod | post | instagram | bubble net feeding |  |
| 13/09/20 | Merimbula (New South Wales) | -36.925544  149.955431 | | | 9:30-12:30 | 100 | post | instagram | lunge feeding |  |
| 16/09/20 | Eden (NSW) | -37.031007  149.982914 | | | Morning | 3 | post | Facebook | Bubble net feeding | \ |
| 17/09/20 | Merimbula (New South Wales) | -36.906878  149.966418 | | | 9:30-12:30 | 2-..-5 | photoes | instagram | lunge feeding |  |
| 17/09/20 | Two Fold Bay | -37.073135  149.929764 | | |  | 1 | description | Facebook | Lunge feeding | \ |
| 18/09/20 | Merimbula (New South Wales) | -36.927170  149.976997 | | | 9:30-12:30 | 2 | \ | instagram | lunge feeding |  |
| 01/10/20 | Barrack Point | -34.561153  150.877514 | | |  | 1 | video | Facebook | \ | \ |
| 06/10/20 | Schouten Island | -42.335308  148.211342 | | |  | 10/20 | videos | facebook | \ | baitfish |
| 07/10/20 | Wineglass Bay | -42.170926  148.305740 | | |  | 12+ | post | facebook | lunge feeding |  |
| 9-10/10/2020 | Jervis Bays | -35.074235  150.754338 | | |  | 150 | post | Facebook | \ | \ |
| 10/10/20 | Jervis Bays | -35.085692  150.744404 | | |  | 2 | post | Facebook | Lunge feeding+ vertical lunge feeding | krill |
| 11/10/20 | Jervis Bays | -35.063154  150.748426 | | |  | 2 | photo | Facebook | Lunge feeding | \ |
| 11/10/20 | Binalong/Bay of Fires | -41.242487  148.354895 | | | afternoon | 6 | video | facebook | lunge feeding |  |
| 11/10/20 | Jervis Bay (NSW) | -35.077018  150.729390 | | |  | 50+ | post | Illawarra Mercury | lunge feeding |  |
| 12/10/20 | Cape Pillar | -43.196761  148.012613 | | |  | 1 | post | facebook | lunge feeding |  |
| 18/10/20 | Tasman Island | -43.229085  148.007850 | | | afternoon | 12+ | **videos** | facebook | lunge feeding |  |
| 18/10/20 | Wilsons Prom | -39.168712  146.398507 | | |  | 2 | videos | Facebook | \ | \ |
| 20/10/20 | Cape Pillar | -43.186616  147.992798 | | |  | 4 | video | facebook | \ |  |
| 21/10/20 | Tasman Peninsula | -43.261650  147.854691 | | |  | 2 | photoes | facebook | \ |  |
| 21/10/20 | Sydney | -33.973458  151.343109 | | | morning | 1 | photoes | instagram | lunge feeding |  |
| 22/10/20 | Tasmania | -43.127468  148.056948 | | | afternoon | 12+ | \ | facebook | \ |  |
| 22/10/20 | south off Hippolyte | -43.123952  148.055306 | | |  | 2 | \ | facebook | \ |  |
| 25/10/20 | Tasmania | -43.022066  147.983847 | | |  | 4? | \ | facebook | lunge feeding |  |
| 31/10/20 | EagleHawk Neck (off Cape Surville) | -42.953045  148.014779 | | | morning | 1+ | \ | facebook | lunge feeding |  |
| 31/10/20 | between Hippolyte and Tasman | -43.125611  148.013149 | | |  | 1+ | \ | facebook | \ |  |
| 08/11/20 | Waterfall bay | -43.059560  147.947881 | | |  | 1+ | \ | facebook | \ | baitfish |
| 08/11/20 | Cape Pillar | -43.202062  147.999693 | | |  | 2-3 | \ | facebook | \ |  |
| 08/11/20 | Tasman Island | -43.234276  147.991198 | | | morning | 1+ | \ | facebook | lunge feeding | krill |
| 15/11/20 | EagleHawk Neck | -43.010714  147.970674 | | | morning | 1+ | \ | facebook | \ |  |
| 15/11/20 | Tasman Peninsula just south of pirates bay Eagle Hawk Neck | -43  148 | | | \ | 30 | \ | Happywhale | lunge feeding + bubbles |  |
| 17/11/20 | between Eaglehawk Neck and Tasman Island | -43.115000  148.013707 | | |  | 2 | \ | facebook | vertical lunge feeding |  |
| 18/11/20 | EagleHawk Neck | -43.023786  147.941504 | | |  | 1+ | \ | facebook | \ |  |
| 18/11/20 | Eaglehawk Neck, Tasman Peninsula | -43.02295168  147.93782942 | | | \ | 20 | \ | Happywhale |  |  |
| 27/11/20 | Pirates Bay | -43.019903  147.934563 | | |  | 1+ | Video | facebook | lunge feeding |  |
| 01/12/20 | Tasman Island | -43.233664  147.994221 | | |  | 1 | \ | facebook | \ |  |
| 21/05/21 | Sydney | -34.063110  151.213127 | | | morning | 1 | \ | instagram | lunge feeding |  |
| 07/06/21 | Wilsons Promontory | -39.087129  146.278316 | | |  | 1 | Video | Facebook | Lunge feeding | Baitfish |
| 8/06/  21 | Wilsons Promontory | -38.994879  146.083951 | | |  | 1 | \ | facebook | lunge feeding | baitfish |
| 11/06/21 | Cape Shanck | -38.499572  144.820918 | | |  | 2 | \ | Facebook | \ | \ |
| 23/07/21 | Killy (Phillip island) | -38.554087  145.379302 | | |  | 1 | \ | Facebook | \ | \ |
| 02/08/21 | Bermagui, New South Wales | -36.448744  150.092547 | | | 10:00 | 2-3 | \ | instagram | bubble net feeding + lunge feeding |  |
| 04/08/21 | Bondi Beach | -33.898814  151.283357 | | | morning | 1 | \ | instagram | lunge feeding |  |
| 07/08/21 | Bermagui, New South Wales | -36.466969  150.092547 | | | 10:00 | 3 | \ | instagram | lunge feeding |  |
| 13/08/21 | Forresters Beach | -33.415478  151.475711 | | |  | 3 | \ | instagram | lunge feeding | baitfish |
| 15/08/21 | Sydney | -33.844822  151.326796 | | | Sunset | 1 | \ | Instagram | Lunge feeding | \ |
| 18/08/21 | Bermagui, New South Wales | -36.441288  150.105936 | | | 10:00 | 1 | \ | instagram | bubble net feeding or vertical lunge feeding |  |
| 19/08/21 | Sapphire Coast | -36.947671  150.004567 | | |  | 1 | \ | instagram | lunge feeding | baitfish |
| 22/08/2021 | Forest caves | -38.527635  145.300579 | | |  | 1 | \ | facebook | \ |  |
| 22/08/21 | Bass Point Memorial (Shellharbour) | -34.595227  150.907822 | | |  | 2 | Photos | Facebook | Lunge feeding | \ |
| 22/08/21 | Surf Beach | -35.761395  150.239536 | | | 9:50 AM | 2 (maybe also 3-4) | \ | facebook | lunge feeding |  |
| 22/08/21 | Surf Beach | -35.761221  150.235206 | | | sunset | 4-…-6 | \ | facebook | lunge feeding | bait ball |
| 30/08/21 | Bermegui | -36.429162  150.092807 | | |  | 1 | \ | Facebook | Lunge feeding | \ |
| 02/09/21 | Bermagui | -36.433858  150.084911 | | |  | megapod | \ | Facebook | Lunge feeding | bait ball |
| 6-12/09/2021 | Bermagui, New South Wales | -36.456753  150.101130 | | | 10:00 | megapod | \ | facebook | lunge feeding | baitfish |
| 09/09/21 | Bermagui, New South Wales | -36.440735  150.084650 | | |  | 100+ | \ | CNN travel | \ | baitfish |
| 9-10/09/2021 | Bermagui, New South Wales | -36.427201  150.098383 | | | afternoon | 150 | \ | facebook | lunge feeding | small bait fish |
| 11/09/21 | Bermagui, New South Wales | -36.407861  150.086710 | | | afternoon | 40-50 | \ | instagram |  |  |
| 11/09/21 | Bermagui, New South Wales | -36.422642  150.104906 | | |  | 150 | Photos | instagram | lunge feeding |  |
| 12/09/21 | Bermagui, New South Wales | -36.407585  150.092718 | | | 10:00 | 80 | \ | instagram | lunge feeding | baitfish |
| 12/09/21 | Bermagui, New South Wales | -36.425129  150.125162 | | |  | 150 | \ | 7 news | lunge feeding | small bait fish |
| 13/09/21 | Bermagui, New South Wales | -36.441978  150.128596 | | | 10:00 | 100+ | \ | instagram | lunge feeding |  |
| 13/09/21 | Port Mcquarie | -31.415226  152.932504 | | |  | 100+ | \ | Newspaper | \ | bait ball |
| 20/09/21 | Peggy's point (Bicheno) | -41.864726  148.305276 | | |  | 2-3 | \ | facebook | \ |  |
| 22/09/21 | Derwent (between Tranmere and Sandy Bay) | -42.910773  147.378050 | | |  | 1 | \ | facebook | lunge feeding | school of fish (salmon) |
| 22/09/2021 | Cape Wellington | -39.067006  146.487939 | | |  | 6 | \ | facebook | \ |  |
| 27/09/21 | Wilsons Prom | -39.044681  146.501106 | | |  | 1+ | Video | Facebook | \ | baitfish |
| 28/09/21 | Tranmere (very end of Oceana Drive) | -42.928320  147.407253 | | |  | 1 |  | facebook | \ |  |
| 06/10/21 | Waterfall bay | -43.058769  147.949661 | | |  | 1 |  | facebook | \ |  |
| 09/10/21 | East North East of pirates bay Tasman peninsula Tasmania 3 miles from shore, Australia | -43.024519  147.993950 | | |  | 20+ | post | happywhale | lunge feeding + bubbles netting | \ |
| 09/10/21 | North of Fortescue Bay, Tasman peninsula, Tasmania, 1 NM from shore | -43.07930377  147.96644210 | | | \ | 2 | post | Happywhale | Lunge feeding + bubble nets feeding |  |
| 09/10/21 | Cosy/swimcart (Binalong Bay) | -41.224422  148.290060 | | |  | 1+ | Photos | facebook | \ |  |
| 12/10/21 | Hippolyte | -43.121294  148.055080 | | |  | 3 | video | facebook | lunge feeding |  |
| 12/10/21 | Hippolyte | -43.120431  148.053970 | | | after dawn | 12 | photo | facebook | lunge feeding |  |
| 12/10/21 | Devenport (12 km off) | -40.655120  146.372249 | | |  | 5-6 | post | facebook | \ |  |
| 12/11/21 | North Cape, Kangaroo Island | -35.555017  137.644567 | | | \ | 20 | Post | Happywhale |  |  |
| 16/11/21 | Western River Cove, South Australia | -35.6685  136.958833 | | | \ | 4 | Photo | Happywhale |  |  |
| 24/11/21 | Southern Spencer Gulf, Australia | -35.5994  137.02675 | | | \ | 3 | **photo** | Happywhale | Lunge feeding | baitball |
| 03/12/21 | Kangaroo Island (off the north Coast) | -35.414156  137.352618 | | |  | 8-9 | post | Article on facebook (sightning by RAD KI: "Research and Discovery Coastal Tours) | lunge feeding |  |
| 04/01/22 | Adventure Bay | -43.338099  147.339513 | | |  | 1 | comment | facebook | \ |  |
| 11/01/22 | Bruny Island (south east) | -43.607199  147.427961 | | | afternoon | 3 | post | facebook | \ |  |
| 14/01/22 | Bruny Island | -43.318993  147.433607 | | |  | 2 | post | facebook | \ |  |
| 15/05/2022 | Cape Woolamai (4-5 km south west --> off Arafura Pearl) | -38.589354  145.283408 | | | 4:30 PM | 1 | Post | facebook | \ |  |
| 04/06/22 | Jervis Bays | -35.098673  150.776232 | | |  | 1 | post | Facebook | Lunge feeding | baitfish |
| 02/07/22 | Cape Wollamai | -38.577679  145.349645 | | |  | 2+ | post | Facebook | \ | \ |
| 04/07/22 | Phillip Island | -38.530935  145.242950 | | |  | 1 | Photo | Facebook | Lunge feeding | Baitfish |
| 06/07/22 | Merimbula (New South Wales) | -36.923033  149.921010 | | | 9:30-12:30 | 5 | **photoes** | instagram | bubble net feeding or vertical lunge feeding? | small bait fish |
| 15/07/22 | Merimbula (New South Wales) | -36.916448  150.022752 | | | 9:30-12:30 | 1 | photoes | instagram | lunge feeding | baitfish |
| 21/07/2022 | Port Welshpool | -38.761281  146.388157 | | |  | 2 | Photos | facebook | lunge feeding | Baitfish |
| 21/07/2022 | Wilson Promontory | -39.180647  146.357352 | | |  | 2 | photoes | facebook | lunge feeding | Baitfish |
| 21/07/22 | Eden (New South Wales) | -37.009933  149.956083 | | |  | 1 | post | instagram + Facebook | vertical lunge feeding | baitfish |
| 22/07/22 | Wilsons Prom | -38.982278  146.484197 | | |  | 2 | photoes | Facebook | Lunge feeding | Baitfish |
| 25/07/22 | Sapphire Coast | -36.814756  150.047139 | | |  | 2 | post | instagram | lunge feeding | baitfish |
| 2/08/ 22 | San Remo (Bore Beach, Potters Hill Road) | -38.542774  145.387334 | | |  | 4 | photoes | facebook | lunge feeding | Baitfish |
| 2/08/ 22 | Bore Beach (San Remo) | -38.539104  145.384913 | | |  | 3-4 | post | facebook | \ |  |
| 3/08/ 22 | Anzacs | -38.546288  145.321444 | | | evening | 2 | photoes | facebook | \ | Baitfish |
| 4/08/ 22 | Cape Woolamai | -38.577262  145.356441 | | | 11:28 AM | 2 | photoes | facebook | \ | school of fish |
| 4/08/ 22 | surfies Point (Philip Island) | -38.517257  145.279935 | | |  | 4-…-7 | video | facebook | lunge feeding | school of fish |
| 5/08/ 22 | Pyramid rock (1st lookout) | -38.527263  145.230663 | | | 12:30 | 1 | post | facebook | \ | school of fish |
| 6/08/ 22 | Pyramid Rock | -38.532890  145.228950 | | | 12:30 | 1 | Photo | facebook | lunge feeding | school of fish |
| 8/08/ 22 | Cape Schanck | -38.531405  144.880442 | | | 2-4 pm | 2 | post | facebook | lunge feeding | school of fish |
| 09/08/22 | Cape Nelson, western Victoria | -38.434983  141.538446 | | | \ | 1 | post | Happywhale |  |  |
| 12/08/22 | Harvey Bay (East coast of Fraser Island) | -25.096168  153.001822 | | |  | 3 | post | facebook | bubble net feeding | baitfish |
| 19/08/2022 | Wilsons Promontory | -38.972553  146.151294 | | |  | 1+ | post | instagram | vertical lunge feeding | baitfish |
| 21/08/22 | Bournda (NSW) | -36.828300  149.959535 | | |  | 1 | post | instagram | lunge feeding | fish bait |
| 22/08/22 | Bermagui, New South Wales | 36.432725  150.108340 | | | afternoon | 1+ | post | instagram | lunge feeding |  |
| 28/08/2022 | South east of Seal Rocks | -38.531799  145.105384 | | |  | 1 | Post | facebook | \ |  |
| 29/08/22 | Wilsons Prom | -38.953247  146.477857 | | |  | 1 | Photo | Facebook | Lunge feeding | \ |
| 29/08/22 | Merimbula (New South Wales) | -36.936772  150.042728 | | | 10:30-12:30 | 1 | Post | instagram | lunge feeding |  |
| 29/08/22 | Bournda (NSW) | -36.760973  150.050923 | | |  | 1 | post | Instagram | Lunge feeding | bait ball |
| 30/08/22 | Twofold Bay (NSW) | -37.071264  149.945933 | | |  | 1 | post | Facebook | Lunge feeding | baitball |
| 7/09/2022 | Seal Rocks | -38.521639  145.098766 | | |  | 5-6 | comment | facebook | \ |  |
| 7/09/2022 | The Nobbies (Phillip Island) | -38.513349  145.111400 | | |  | 3 | Photos | facebook | \ |  |
| 9/09/2022 | Kilicunda (off the saddleback) | -38.590690  145.502086 | | |  | 2 | Post | facebook | \ |  |
| 12/09/22 | Twofold Bay (NSW) | -37.087364  149.913900 |  | | | 1 | Post | Facebook | Lunge feeding | baitball |
| 13/09/22 | Twofold Bay (NSW) | -37.061295  149.930557 | | |  | 1 | Video | Facebook | Lunge feeding | baitball |
| 13/09/22 | Ulverstone (15 kms off) | -40.711158  146.245960 | | |  | 4 | Post | facebook | vertical lunge feeding | baitfish |
| 14/09/2022 | Shelleys Beach (Kilcunda) | -38.557707  145.465534 | | | 2:45 PM | 1 | Post | facebook (Phillip Island Whale Watchers) | \ |  |
| 14/09/2022 | Shelleys Beach (Kilcunda) | -38.555777  145.468770 | | | 4:00 PM | 1 | Video | facebook | \ |  |
| 14/09/22 | Twofold Bay (NSW) | -37.075353  149.959067 |  | | | 2 |  | Sarah McCulloch |  |  |
| 15/09/22 | Twofold Bay (NSW) | -37.082764  149.911017 | | |  | 1 |  | Facebook | Lunge feeding + vertical lunge feeding | \ |
| 17/09/22 | Hobart/Nipaluna | -42.893181  147.348119 | | |  | 1 |  | instagram | vertical lunge feeding |  |
| 17/09/22 | Iron pot (south of) | -43.062009  147.416931 | | | afternoon | 3 (but just 1 feeding) | Photo | facebook | \ | baitfish |
| 19/09/22 | K'Gari (Fraser Island) | -24.898478  153.159160 | | |  | 2+ | Post | Facebook | \ | \ |
| 20/09/22 | Merimbula (New South Wales) | -36.884014  150.016698 | | | 9:30-12:30 | 1 | photoes | instagram | lunge feeding |  |
| 22/09/2022 | Kilcunda | -38.567213  145.465137 | | | 1-2 pm | 2 | photoes | facebook | \ | baitfish |
| 01/10/22 | south of the Iron Pot | -43.062636  147.416931 | | | morning | 1 | **photos** | facebook | \ | Atlantic salmon |
| 02/10/22 | EagleHawk Neck | -43.016219  147.947621 | | |  | many (6) | **photoes** | facebook | \ |  |
| 03/10/22 | Eaglehawk Neck (7 kms E) | -43.027225  148.069944 | | |  | 3 | photoes | facebook | vertical lunge feeding |  |
| 09/10/22 | Merimbula (New South Wales) | -36.906767  149.962218 | | | 9:30-12:30 | 1 | Video | instagram | lunge feeding |  |
| 16/10/22 | East Coast of Tasman Peninsula | -43.101606  148.147361 | | |  | 3 | video | facebook | lunge feeding |  |
| 19/10/22 | Tasman Island | -43.239619  147.990143 | | | morning | 12 | Photos | facebook | \ |  |
| 20/10/22 | Tasman Peninsula | -43.128543  147.597739 | | |  | 1 | photo | facebook | vertical lunge feeding |  |
| 04/11/22 | Western River, Australia | -35.67595  136.922917 | | | \ | 1 | photoes | Happywhale | lunge feeding |  |
| 04/11/22 | off the coast of Kangaroo Island, Australia | -35.668067  136.929067 | | | \ | 1 | photoes | Happywhale |  |  |
| 26/12/22 | Tasman Island | -43.239255  148.017995 | | |  | 1 | photo | facebook | \ |  |
| 24/05/23 | Bass Point | -34.597380  150.905843 | | | 12:20 | 2 | video | Facebook | \ | \ |
| 30/05/23 | Kiama Heights | -34.706173  150.854647 | | | 1-3 pm | 4 | Video | Facebook | Lunge feeding | \ |
| 09/06/23 | Byron Bay | -28.622008  153.619808 | | |  | 1 | Photos | Facebook | Lunge feeding | \ |
| 09/06/23 | Shellharbour | -34.574127  150.875241 | | |  | 1 | **photoes** | Facebook | Lunge feeding | \ |
| 13/06/23 | Bushrangers Bay | -34.597846  150.902011 | | |  | 5 | **article (news)** | Facebook | \ | \ |
| 24/06/23 | mouth of powlett | -38.589553  145.475596 | | | 13:00 | 2-3 | **video** | Facebook | \ | \ |
| 14/07/23 | Port Stephens (NSW) | -32.778840  152.152607 | | |  | 1 | description | Facebook | Lunge feeding | school of fish |
| 27/07/23 | Rhyll | -38.476556  145.314534 | | | 08:30 | 3 | photoes | Facebook | vertical lunge feeding | Baitfish |
| 27/07/23 | Sydney | -33.865350  151.308943 | | |  | 1 | video | Facebook | Lunge feeding | \ |
| 06/08/23 | Bournda (NSW) - 3/4 km from the shore | -36.835653  149.976545 | | |  | 30-40 | **video** | Facebook | Lunge feeding | \ |
| 07/08/23 | Bournda (NSW) | -36.783805  150.098035 | | |  | 2+ | **video** | Instagram | Lunge feeding | \ |
| 07/08/23 | Bournda (NSW) | -36.783805  150.098035 | | |  | 8 | photoes | Instagram | Lunge feeding | \ |
| 15/08/23 | Bournda (NSW) | -36.842078  150.056646 | | |  | 40+ | photo | Instagram | Lunge feeding + bubble net feeding | \ |
| 16/08/23 | Eden (NSW) | -37.083961  149.970564 | | |  | 2 | **photo** | Instagram | Lunge feeding | \ |
| 22/08/23 | Twofold Bay (NSW) | -37.095236  149.914244 | | |  | 1+ | video | Instagram | Lunge feeding | \ |
| 25/08/23 | Eden (NSW) | -37.093618  149.977737 | | | Morning | 1+ | **video** | Instagram | Lunge feeding | \ |
| 21/09/23 | Tasmania, Australia | -42.138252  148.363266 | | | \ | 20-25 | **photo** | Happywhale |  | krill |
| 23/09/23 | Tasmania, Australia | -43.184067  147.987983 | | | \ | 1 | photo | Happywhale |  |  |
| 25/09/23 | Gates Bluff, Coles Bay, Tasmania | -42.228986  148.3461 | | | \ | 15-20 | video | Happywhale |  |  |
| 27/09/23 | Tasmania, Australia | -43.201814  148.003962 | | | \ | 5 | Photo | Happywhale |  | baitfish |
| 27/09/23 | Tasmania, Australia | -43.128587  148.004618 | | | \ | 10-12 | Photos | Happywhale |  |  |
| 30/09/23 | Tasman Island | -41.810711  148.506895 | | |  | 2 | Photo | Facebook | Lunge feeding | \ |
| 06/10/23 | Bass Point | -34.594180  150.904783 | | |  | 1 | photoes | Facebook | Lunge feeding | \ |
| 08/10/23 | Tasmania, Australia | -43.016976  148.000946 | | | \ | 1 | Photo | Happywhale |  |  |
| 08/10/23 | Tasmania, Australia | -43.016976  148.000946 | | | \ | 3 | photo | Happywhale | Lunge feeding |  |
| 09/10/23 | Tasmania, Australia | -43.0292  147.993983 | | | \ | 2 | photo | Happywhale | Lunge feeding |  |
| 10/10/23 | Tasmania, Australia | -43.084346  148.016699 | | | \ | 6 | Post - deriva da post Instagram | Happywhale | Lunge feeding | zooplankton |
| 30/10/23 | Tasmania, Australia | -43.127631  148.023158 | | | \ | 1 | video | Happywhale |  |  |
| 04/12/23 | Boreel Head | -43.515095  147.305756 | | | \ | 1 | video | Happywhale |  |  |
| 23/05/24 | Cape Woolamai pinnacles (500 m from the cape) | -38.568461  145.360042 | | |  | 1-2 | video | Facebook | \ | \ |
| 25/05/24 | Two Bays | -38.315531  144.894499 | | |  | 1 | Photos | Facebook | \ | \ |
| 26/05/24 | Two Bays | -38.289128  144.927458 | | |  | 1 | Photos | Facebook | \ | \ |
| 29/05/24 | Cape Pillar | -43.183004  148.003442 | | | \ | 1 | Photos | Happywhale |  | baitfish |
| 29/05/24 | Shellharbour | -34.577904  150.876817 | | | Afternoon | 3 | Photos | Facebook | \ | \ |
| 08/06/24 | couple of kms out from mouth of powlett. | -38.594438  145.483052 | | | 13:40 | 1 | Photos | Facebook | \ | \ |
| 19/06/24 | San Remo | -38.549580  145.395049 | | |  | 1+ | Photos | Facebook | Lunge feeding | \ |
| 29/06/24 | Tasmania, Australia | -43.243215  147.992334 | | | \ | 2 | Photos | Happywhale |  |  |
| 06/07/24 | Phillip Island | -38.522877  145.271790 | | |  | 2+ | Photos | Facebook | \ | baitfish |
| 10/07/24 | Tasman Peninsula | -42.991482  147.782706 | | |  | 2 | Photos | Facebook | Lunge feeding | \ |
| 27/07/24 | Shellharbour | -34.572651  150.869937 | | |  | 1 |  | Facebook | Lunge feeding | \ |
| 29/07/24 | Shellharbour | -34.572120  150.874595 | | |  | 1 |  | Facebook | Lunge feeding | \ |
| 10/08/24 | Byron Bay | -28.615830  153.601269 | | | evening | 1+ |  | Facebook | Lunge feeding | \ |
| 15/08/24 | Tasmania, Australia | -43.068802  147.392881 | | | \ | 1 | photoes | Happywhale | Lunge feeding |  |
| 18/08/24 | Tasmania, Australia | -43.066124  147.384253 | | | \ | 1 | Photos | Happywhale |  |  |
| 20/08/24 | Wilsons Prom | -39.089732  146.457946 | | |  | 1 | video | Facebook | \ | \ |
| 22/08/24 | Bruny Island | -43.546704  147.432790 | | |  | 1 | Photo | Facebook | Vertical lunge feeding | \ |
| 23/08/24 | Platypus Bay | -24.952561  153.09143 | | | \ | 1 | Photos | Happywhale |  |  |
| 08/09/24 | Merimbula (NSW) | -36.873904  149.941619 | | | Afternoon | 8 | Photos | Instagram | \ | \ |
| 08/09/24 | Twofold (NSW) | -37.080986  149.951921 | | | Morning | 1 | Photos | Instagram | Lunge feeding | baitfish |
| 08/09/24 | Merimbula bay (NSW) | -36.910114  149.923015 | | | Morning (till afternoon) | 2+ | Video | Instagram | \ | \ |
| 09/09/24 | Tasmania, Australia | -43.18649  148.007653 | | | \ | 4 | Post | Happywhale |  |  |
| 09/09/24 | Merimbula (NSW) | -36.878023  149.942048 | | | Morning | 8+ | photoes | Instagram | Lunge feeding | \ |
| 11/09/24 | Merimbula (NSW) | -36.874384  149.956897 | | |  | 1 | photoes | Instagram | Lunge feeding | \ |
| 14/09/24 | Byron Bay | -28.616263 153.606043 | |  | | 1 | Post | Instagram | \ | \ |
| 15/09/24 | Masons Cove and NW Port Arthur | -43.150263  147.864037 | | | \ | 1 | Photos | Happywhale | Lunge feeding | baitfish |
| 16/09/24 | Western River Cove | -35.666013  136.97199 | | | \ | 3 | Photo | Happywhale | Lunge feeding | anchovies |
| 17/09/24 | Shellharbour | -34.590670  150.893730 | | | 12:00 | 1 | photoes | Facebook | Lunge feeding + vertical lunge feeding | baitfish |
| 20/09/24 | Leonards Island (NSW) | -37.017283  149.948681 | | |  | 4 | photo | Instagram | Lunge feeding | \ |
| 21/09/24 | Byron Bay | -28.659933  153.639103 | | |  | 1 | photo | Instagram | Lunge feeding | \ |
| 22/09/24 | Bass Point | -34.595868  150.904429 | | |  | 1 | photo | Facebook | Lunge feeding | \ |
| 23/09/24 | The Thumbs | -43.10489  147.98333 | | | \ | 2 | **photo** | Happywhale |  |  |
| 24/09/24 | Western River Cove | -35.664002  136.971446 | | | \ | 4 | **video** | Happywhale |  | anchovies |
| 26/09/24 | Snellings Beach, Kangaroo Island | -35.658444  137.063048 | | | \ | 2 | video | Happywhale | Lunge feeding | anchovies |
| 28/09/24 | Phillip Island | -38.527983  145.177155 | | |  | 5 | post | Facebook | \ | \ |
| 04/10/24 | Yanky Rock, Black Shore | -43.223478  147.980363 | | | \ | 1 | photoes | Happywhale |  | baitfish |
| 04/10/24 | Wilsons Prom | -39.032880  146.500270 | | |  | 2 | article (news) | Facebook | Lunge feeding? | \ |
| 06/10/24 | Mount Brown | -43.213179  147.876921 | | | \ | 2 | post | Happywhale |  |  |
| 09/10/24 | Cape Wollamai (2km SW) | -38.582972  145.334508 | | | 11:30 | 2 | post | Facebook | \ | \ |
| 09/10/24 | Ulverstone | -41.118655  146.198419 | | | \ | 2 | video | Happywhale |  | baitfish |
| 15/10/24 | Near Betsy Island | -43.087334  147.537328 | | | \ | 1 | \ | Happywhale |  |  |
| 16/10/24 | near Black Heads | -43.215749  147.946429 | | | \ | 1 | \ | Happywhale |  |  |
| 20/10/24 | Hippolyte Rock | -43.117361  148.054634 | | | \ | 1 | \ | Happywhale |  |  |
| 23/10/24 | Port Welshpool | -38.755229  146.388287 | | |  | 4 | \ | Facebook | \ | \ |
| 24/10/24 | East of Wineglass Bay, Australia | -42.159178  148.412709 | | | \ | 20-30 | article (news) | Happywhale |  |  |
| 10/11/24 | NE of Cape Hauy | -43.089035  147.996106 | | | \ | 3 | Photos | Happywhale |  |  |
| 13/11/24 | West of Tasman | -43.241873  148.013141 | | | \ | 2 | \ | Happywhale | Lunge feeding |  |
| 13/11/24 | Off Ulverstone | -41.122645  146.17996 | | | \ | 2 | photoes | Happywhale |  |  |
| 14/11/24 | Eagleawk Neck | -43.043301  148.040388 | | |  | 1 | **photoes** | Facebook | Vertical lunge feeding | baitfish |
| 14/11/24 | Smith Bay, Kangaroo Island | -35.555444  137.447287 | | | \ | 1 | photoes | Happywhale | Lunge feeding | krill |
| 16/11/24 | East of Tasman Island | -43.239685  148.01895 | | | \ | 2 | \ | Happywhale |  |  |
| 16/11/24 | Black Head | -43.215749  147.946429 | | | \ | 3 | \ | Happywhale |  |  |
| 17/11/24 | Black Head | -43.215749  147.946429 | | | \ | 3 | Post | Happywhale |  |  |
| 21/11/24 | Cape Hauy | -43.130786  148.012659 | | | \ | 3 | Photo | Happywhale | Lunge feeding |  |
| 21/11/24 | Munroe Bight | -43.183487  148.007286 | | | \ | 2 | video | Happywhale | Lunge feeding |  |
| 23/11/24 | Hippolyte Rock | -43.117248  148.048047 | | | \ | 1 | Video | Happywhale |  |  |
| 24/11/24 | Cape Hauy | -43.131796  148.005712 | | | \ | 2 | Video | Happywhale |  |  |
| 24/11/24 | Fortescue Bay | -43.122923  147.991732 | | | \ | 1 | Photos | Happywhale |  |  |
| 25/11/24 | Eagleawk Neck | -43.021440  147.991702 | | |  | 2 | Photo | Facebook | Lunge feeding | \ |
| 25/11/24 | Mouth of Fortescue Bay | -43.129696  147.984385 | | | \ | 8-10 | Photos | Happywhale |  |  |
| 26/11/24 | Between the Hippolyte Rock and the Thumbs | -43.111904  148.019748 | | | \ | 2 | Photo | Happywhale |  |  |
| 27/11/24 | East of Tasman | -43.239485  148.017263 | | | \ | 1 | Post | Happywhale |  |  |
| 28/11/24 | Tasman Island | -41.181530  148.531244 | | |  | 3 | Video | Facebook | Vertical lunge feeding + Lunge feeding | \ |
| 29/11/24 | Munro Bight | -43.176533  148.007388 | | | \ | 1 | Video | Happywhale |  |  |
| 05/12/24 | Emu Bay, Kangaroo Island | -35.563473  137.522979 | | | \ | 1 | Photos | Happywhale | Lunge feeding | krill |
| 08/12/24 | Munroe Bight | -43.185408  147.992284 | | | \ | 1 | \ | Happywhale |  |  |
| 14/12/24 | Munroe Bight | -43.185938  147.990375 | | | \ | 1 | Photo | Happywhale |  | baitball |
| 08/01/25 | West off South Island Bruny | -43.418297  147.387032 | | | \ | 1 | \ | Happywhale |  |  |
| 16/01/25 | Bruny Island | -43.398185  147.431717 | | |  | 1 |  | Facebook | Lunge feeding | baitfish |

TABLE S2.

| **Operator** | **State(s)** | **Region(s) served** | **Departure port(s)** | **Active during study period** | **Source** |
| --- | --- | --- | --- | --- | --- |
| Sea World Cruises | QLD | Gold Coast | Main Beach (Gold Coast) | 2020–2025 | [https://seaworldcruises.com.au](https://seaworldcruises.com.au/) |
| Whales in Paradise | QLD | Gold Coast | Surfers Paradise; Marina Mirage | 2020–2025 | [https://www.whalesinparadise.com.au](https://www.whalesinparadise.com.au/) |
| Spirit of Gold Coast | QLD | Gold Coast | Main Beach (Gold Coast) | 2020–2025 | [https://www.spiritofgoldcoast.com.au](https://www.spiritofgoldcoast.com.au/) |
| Whale Watch Queensland | QLD | Gold Coast | Surfers Paradise; Sanctuary Cove | 2020–2025 | [https://whalewatchqueensland.com](https://whalewatchqueensland.com/) |
| Cooly Eco Adventures | QLD | Southern Gold Coast | Coolangatta | 2020–2025 | [https://coolyecoadventures.com.au](https://coolyecoadventures.com.au/) |
| Kokomo Cruises | QLD | Gold Coast | Gold Coast | 2020–2025 | [https://kokomocruises.com.au](https://kokomocruises.com.au/) |
| Brisbane Whale Watching | QLD | Brisbane; Moreton Bay | Redcliffe | 2020–2025 | [https://brisbanewhalewatching.com.au](https://brisbanewhalewatching.com.au/) |
| Tangalooma Whale Watching Cruises | QLD | Moreton Bay | Brisbane; Tangalooma (Moreton Island) | 2020–2025 | [https://www.tangalooma.com](https://www.tangalooma.com/) |
| Sunreef Mooloolaba | QLD | Sunshine Coast | Mooloolaba | 2020–2025 | [https://sunreef.com.au](https://sunreef.com.au/) |
| Whale One | QLD | Sunshine Coast | Mooloolaba | 2020–2025 | [https://whaleone.com.au](https://whaleone.com.au/) |
| Epic Ocean Adventures | QLD | Sunshine Coast | Mooloolaba | 2020–2025 | [https://epicoceanadventures.com.au](https://epicoceanadventures.com.au/) |
| Blue Dolphin Marine Tours | QLD | Fraser Coast / Hervey Bay | Urangan Harbour (Hervey Bay) | 2020–2025 | [https://bluedolphin.com.au](https://bluedolphin.com.au/) |
| Tasman Venture | QLD | Fraser Coast / Hervey Bay | Great Sandy Straits Marina (Hervey Bay) | 2020–2025 | [https://tasmanventure.com.au](https://tasmanventure.com.au/) |
| Spirit of Hervey Bay | QLD | Fraser Coast / Hervey Bay | Great Sandy Straits Marina (Hervey Bay) | 2020–2025 | [https://www.spiritofherveybay.com.au](https://www.spiritofherveybay.com.au/) |
| Whalesong Cruises | QLD | Fraser Coast / Hervey Bay | Urangan Harbour (Hervey Bay) | 2020–2025 | [https://whalesong.com.au](https://whalesong.com.au/) |
| Hervey Bay Whale Watch | QLD | Fraser Coast / Hervey Bay | Urangan Harbour (Hervey Bay) | 2020–2025 | [https://www.herveybaywhalewatch.com.au](https://www.herveybaywhalewatch.com.au/) |
| Freedom Whale Watch | QLD | Fraser Coast / Hervey Bay | Hervey Bay | 2020–2025 | [https://freedomwhalewatch.com.au](https://freedomwhalewatch.com.au/) |
| Hervey Bay Dive Centre | QLD | Fraser Coast / Hervey Bay | Urangan Harbour (Hervey Bay) | 2020–2025 | [https://www.herveybaydivecentre.com.au](https://www.herveybaydivecentre.com.au/) |
| Salty Safaris | QLD | Fraser Coast / Hervey Bay | Great Sandy Straits Marina (Hervey Bay) | 2020–2025 | [https://salty-safaris.com.au](https://salty-safaris.com.au/) |
| Australia Whale Experience / Lady Musgrave Experience | QLD | Bundaberg; Southern Great Barrier Reef | Burnett Heads / Bundaberg Port Marina | 2020–2025 | [https://australiawhaleexperience.com.au](https://australiawhaleexperience.com.au/) |
| Hervey Bay Whale Watch | QLD | Fraser Coast / Hervey Bay / Platypus Bay | Great Sandy Straits Marina, Urangan | 2020–2025 | [https://www.herveybaywhalewatch.com.au](https://www.herveybaywhalewatch.com.au/?utm_source=chatgpt.com) |
| Brisbane Whale Watching | QLD | Moreton Bay / Brisbane | Redcliffe | 2020–2025 | [https://brisbanewhalewatching.com.au](https://brisbanewhalewatching.com.au/?utm_source=chatgpt.com) |
| Tangalooma Whale Watching | QLD | Moreton Island / Moreton Bay | Tangalooma / Brisbane | 2020–2025 | <https://www.tangalooma.com/whale-watching> |
| Byron Bay Whale Watching | NSW | Byron Bay; Northern Rivers | Byron Bay | 2020–2025 | [https://www.byronbaywhalewatching.com.au](https://www.byronbaywhalewatching.com.au/) |
| Out of the Blue Adventures | NSW | Ballina; Northern Rivers | Ballina | 2020–2025 | [https://outoftheblueadventures.com.au](https://outoftheblueadventures.com.au/) |
| Pacific Explorer | NSW | Coffs Coast; Solitary Islands | Coffs Harbour | 2020–2025 | [https://www.pacificexplorer.com.au](https://www.pacificexplorer.com.au/) |
| Jetty Dive | NSW | Coffs Coast; Solitary Islands | Coffs Harbour | 2020–2025 | [https://jettydive.com.au](https://jettydive.com.au/) |
| Port Jet Cruises | NSW | Port Macquarie; Mid North Coast | Port Macquarie | 2020–2025 | [https://www.portjet.com.au](https://www.portjet.com.au/) |
| Moonshadow–TQC Cruises | NSW | Port Stephens | Nelson Bay | 2020–2025 | [https://www.moonshadow.com.au](https://www.moonshadow.com.au/) |
| Imagine Cruises | NSW | Port Stephens | Nelson Bay | 2020–2025 | [https://www.imaginecruises.com.au](https://www.imaginecruises.com.au/) |
| Aquamarine Adventures | NSW | Port Stephens | Nelson Bay | 2020–2025 | [https://www.aquamarineadventures.com.au](https://www.aquamarineadventures.com.au/) |
| CoastXP | NSW | Newcastle; Hunter Coast | Newcastle Harbour | 2020–2025 | [https://coastxp.com.au](https://coastxp.com.au/) |
| Whale Watching Sydney | NSW | Sydney; Sydney coastal waters | Sydney Harbour | 2020–2025 | [https://www.whalewatchingsydney.com.au](https://www.whalewatchingsydney.com.au/) |
| Ocean Extreme | NSW | Sydney; Sydney coastal waters | Sydney Harbour | 2020–2025 | [https://oceanextreme.com.au](https://oceanextreme.com.au/) |
| Fantasea Cruising | NSW | Sydney; Sydney coastal waters | Circular Quay; Darling Harbour | 2020–2025 | [https://fantasea.com.au](https://fantasea.com.au/) |
| Oz Whale Watching | NSW | Sydney; Sydney Harbour; offshore Sydney | Circular Quay; Darling Harbour | 2020–2025 | [https://www.ozwhalewatching.com.au](https://www.ozwhalewatching.com.au/) |
| Jervis Bay Wild | NSW | Jervis Bay; Shoalhaven | Huskisson | 2020–2025 | [https://www.jervisbaywild.com.au](https://www.jervisbaywild.com.au/) |
| Dolphin Watch Cruises Jervis Bay | NSW | Jervis Bay; Shoalhaven | Huskisson | 2020–2025 | [https://www.dolphinwatch.com.au](https://www.dolphinwatch.com.au/) |
| Shellharbour Wild | NSW | Illawarra; Shellharbour | Shellharbour | 2020–2025 | [https://shellharbourwild.com.au](https://shellharbourwild.com.au/) |
| Cat Balou Cruises | NSW | Sapphire Coast; Twofold Bay; Eden | Eden | 2020–2023 | [https://www.catbalou.com.au](https://www.catbalou.com.au/) |
| Sapphire Coastal Adventures | NSW | Sapphire Coast; Twofold Bay; Eden; Merimbula; Bermagui | Eden; Merimbula | 2020–2025 | [https://www.sapphirecoastaladventures.com.au](https://www.sapphirecoastaladventures.com.au/) |
| Go Whale Watching | NSW | Sydney; Sydney coastline | Darling Harbour | 2020–2025 | [https://www.gowhalewatching.com.au](https://www.gowhalewatching.com.au/) |
| NOVA Cruises | NSW | Newcastle; Hunter; Lake Macquarie | Newcastle Harbour | 2020–2025 | [https://novacruises.com.au](https://novacruises.com.au/) |
| Byron Bay Whale Watching | NSW | Byron Bay; Northern Rivers | Byron Bay | 2020–2025 | [https://www.byronbaywhalewatching.com.au](https://www.byronbaywhalewatching.com.au/) |
| Blue Bay Whale Watching | NSW | Byron Bay; Northern Rivers | Byron Bay | 2020–2025 | [https://www.bluebaywhalewatching.com.au](https://www.bluebaywhalewatching.com.au/) |
| Solitary Islands Marine Discoveries | NSW | Coffs Coast; Solitary Islands | Coffs Harbour | 2020–2025 | [https://solitaryislands.com.au](https://solitaryislands.com.au/) |
| Dolphin Watch Cruises | NSW | Jervis Bay; Shoalhaven | Huskisson | 2020–2025 | [https://www.dolphinwatch.com.au](https://www.dolphinwatch.com.au/) |
| Freedom Whale Watch | NSW | Sapphire Coast; Eden | Eden | 2020–2025 | [https://www.freedomwhalewatch.com.au](https://www.freedomwhalewatch.com.au/) |
| Wild Byron | NSW | Byron Bay / Northern Rivers / Cape Byron Marine Park | Brunswick Heads | 2020–2025 | [https://wildbyron.com.au](https://wildbyron.com.au/?utm_source=chatgpt.com) |
| Byron Bay Whale Watching | NSW | Byron Bay | Byron Bay | 2020–2025 | [https://www.byronbaywhalewatching.com.au](https://www.byronbaywhalewatching.com.au/?utm_source=chatgpt.com) |
| Solitary Islands Marine Discoveries | NSW | Coffs Harbour / Solitary Islands Marine Park | Coffs Harbour | Yes* | (official website to verify) |
| Port Jet Cruise Adventures | NSW | Port Macquarie / Mid North Coast | Port Macquarie | 2020–2025 | [https://portjet.com.au](https://portjet.com.au/?utm_source=chatgpt.com) |
| Epic Surf Ocean Adventures | NSW | Forster / Great Lakes | Forster | 2020–2025 | [https://www.forsterluxuryhouseboats.com.au/whale-dolphin-and-lake-tours/](https://www.forsterluxuryhouseboats.com.au/whale-dolphin-and-lake-tours/?utm_source=chatgpt.com) |
| Terrigal Ocean Tours | NSW | Central Coast | Terrigal Haven | 2020–2025 | [https://terrigaltours.com.au](https://terrigaltours.com.au/) |
| Cronulla Whale Watching | NSW | Sydney South / Sutherland Shire | Cronulla | 2020–2025 | [https://www.cronullawhalewatching.com.au](https://www.cronullawhalewatching.com.au/?utm_source=chatgpt.com) |
| Wildlife Coast Cruises | VIC | Phillip Island; Bass Coast; Wilsons Promontory; Gippsland | Cowes; Port Welshpool | 2020–2025 | [https://wildlifecoastcruises.com.au](https://wildlifecoastcruises.com.au/) |
| Wanderer Adventures (Wilsons Promontory Cruises Pty Ltd) | VIC | Wilsons Promontory; Gippsland | Tidal River (Wilsons Promontory) | 2020–2025 | [https://www.wandereradventures.com.au](https://www.wandereradventures.com.au/) |
| Phillip Island Whale Watching | VIC | Phillip Island; Bass Strait | San Remo | 2020–2025 | [https://www.phillipislandwhalewatching.com.au](https://www.phillipislandwhalewatching.com.au/) |
| Portland Ocean Adventures | VIC | Portland; Discovery Bay; western Victoria | Portland | 2020–2025 | [https://portlandoceanadventures.com.au](https://portlandoceanadventures.com.au/) |
| Pennicott Wilderness Journeys | TAS | Bruny Island; Tasman Peninsula; Southeast Tasmania; Freycinet / East Coast Tasmania | Adventure Bay; Hobart; Port Arthur | 2020–2025 | [https://www.pennicottjourneys.com.au](https://www.pennicottjourneys.com.au/) |
| Wild Ocean Tasmania | TAS | Tasman Peninsula; Southern Tasmania | Eaglehawk Neck | 2020–2025 | [https://wildoceantasmania.com.au](https://wildoceantasmania.com.au/) |
| Pirates Bay Charters | TAS | Tasman Peninsula | Eaglehawk Neck | 2020–2025 | <https://www.facebook.com/piratesbaycharters> |
| South East Charters | TAS | Tasman Peninsula; Storm Bay | Eaglehawk Neck | 2020–2025 | [https://www.southeastcharters.com.au](https://www.southeastcharters.com.au/) |

TABLE S3.

| State | Region | Feeding records (n) | Whale-watching operators (n) | Observation effort proxy |
| --- | --- | --- | --- | --- |
| QLD | Fraser Coast (Hervey Bay) | 3 | 9 | Very High |
| QLD | Sunshine Coast | 0 | 3 | Moderate |
| QLD | Gold Coast | 0 | 5 | High |
| NSW | Byron Bay / Northern Rivers | 4 | 6 | High |
| NSW | Port Stephens | 1 | 3 | Moderate |
| NSW | Sydney | 10 | 7 | Very High |
| NSW | Illawarra | 14 | 1 | Low |
| NSW | Jervis Bay / Shoalhaven | 7 | 3 | Moderate |
| NSW | Sapphire Coast | 53 | 3 | Moderate |
| VIC | Phillip Island / Bass Strait | 50 | 3 | Moderate |
| TAS | Southeast Tasmania | 96 | 4 | Moderate |
| SA | South Australia | 11 | 0 | Low |
|  | Other | 7 | 16 | Very High |
|  | **Total** | **256** | **63** | — |
